# Electron transport chain disruption leads to failure of glucose derepression via the Snf1/AMPK pathway

**DOI:** 10.1101/2025.11.06.687039

**Authors:** Alisha G. Lewis, Jeremy R. Smith, Matthew C. Smardz, Celeste M. Sandoval, Sho W. Suzuki, Bryson D. Bennett, Patrick A. Gibney

## Abstract

When *Saccharomyces cerevisiae* cells transition from a glucose-rich environment to low glucose conditions, the expression of genes that were previously repressed by glucose is derepressed, enabling the cell to adapt metabolic processes to the available carbon source. The Snf1 pathway is one of the primary signaling pathways responsible for orchestrating glucose sensing and signaling. In this study, we investigate the impact of disrupted electron transport chain (ETC) function, a mitochondrial protein complex essential for respiratory energy generation, in glucose derepression. We observe that respiratory incompetent mutants exposed to glucose are unable to subsequently utilize galactose as a carbon source in minimal media. In contrast, ETC mutants that have been generated and maintained on galactose can effectively continue to metabolize galactose until glucose exposure. We define this phenomenon as a <u>F</u>ailure of <u>G</u>lucose <u>D</u>erepression (FGD), wherein respiratory incompetent cells fail to fully reverse glucose repressed gene expression regulation. Through further characterization, we show how irregular localization patterns of crucial proteins within the Snf1 pathway are associated with FGD suggesting a potential novel connection between the ETC and the Snf1 pathway during carbon source transition.

**SIGNIFICANCE STATEMENT:**

1. Respiratory incompetent electron-transport chain mutants are capable of metabolizing galactose as a carbon source. However, when ETC mutants encounter glucose, they lose this ability to metabolize galactose.
2. This study demonstrates that failure to derepress glucose repressed genes facilitated by the Snf1/AMPK pathway is primarily responsible for this phenotype in ETC mutants.
3. These results illustrate a novel signaling role for the ETC and suggest a possible metabolic intervention to certain disease phenotypes.

## INTRODUCTION

Glucose, the preferred carbon source of *Saccharomyces cerevisiae*, is predominantly metabolized through fermentation rather than respiration when present at high concentration, including when oxygen is readily available, a phenomenon known as the Crabtree effect in yeast (a similar phenomenon is termed the Warburg effect in cancer cells) (Diaz-Ruiz et al., 2011). High glucose concentrations repress the expression of genes involved in alternative carbon source utilization, respiration, gluconeogenesis, peroxisomal function, and fatty acid metabolism (Carlson, 1999). However, upon glucose depletion, these glucose repressed genes are derepressed, enabling the yeast cell to express relevant genes and optimize energy utilization by metabolizing available carbon sources. While several glucose signaling systems have been identified, including the Ras/PKA and Rgt2/Snf3 pathways, the Snf1 pathway primarily governs cell survival under low glucose conditions or when less preferred alternative carbon sources are present (Carlson, 1999; Hedbacker and Carlson, 2009; Zaman et al., 2009)

In conditions of abundant glucose, growth-promoting pathways such as Ras/PKA and TOR are activated, facilitating cell growth and proliferation (Werner-Washburne et al., 1993; Herman, 2002). However, when glucose becomes limited, and the cell must rely on alternate carbon sources for energy, such as galactose, the Snf1 complex is activated, phosphorylating downstream effectors, and regulating the transcription of glucose-repressed genes. In addition to its established role under glucose-depleted conditions, the Snf1 complex is involved in adapting to various environmental stresses and exhibits crosstalk with other nutrient-responsive pathways (Hong and Carlson, 2007; Shashkova et al., 2015). The Snf1 complex consists of an alpha catalytic subunit (Snf1), one of three beta subunits (Gal83, Sip1, Sip2) and a gamma subunit (Snf4). Activated under glucose derepressing conditions (e.g. low glucose), the Snf1 protein kinase regulates a subset of glucose-repressed genes by directly phosphorylating transcriptional coactivators such as Cat8 and Adr1 and by inhibiting the repressor activity of Mig1 (Randez-Gil et al., 1997; Treitel et al., 1998; Ratnakumar et al., 2009). The high degree of evolutionary conservation between Snf1 and its mammalian equivalent, AMPK, underscores the functional significance of this pathway as a key regulator of energy homeostasis (Hedbacker and Carlson, 2009).

Concomitant with Snf1 activation, the metabolic machinery dedicated to respiratory processes is upregulated, allowing the cell to redirect its metabolism toward mitochondrial function. Through targeting transcriptional activators, the Snf1 kinase regulates the expression of mitochondrial genes, including those involved in the tricarboxylic acid cycle and electron transport chain (ETC) (Deng et al., 2023). The ETC, a series of protein complexes located in the inner mitochondrial membrane, primarily functions in ATP generation through oxidative phosphorylation. While the ETC has traditionally been viewed as a metabolic pathway, previous publications have unveiled novel signaling roles associated with this protein complex. In mammalian cells for instance, cytochrome *c*, an ETC protein involved in electron transfer, also initiates apoptosis (Cai et al., 1998). Upon release from the mitochondrial membrane, cytochrome *c* allosterically activates apoptosis-protease activating factor 1 in the presence of ATP, triggering a cascade of caspase reactions that ultimately lead to cell death (Garrido et al., 2006). Similarly, other mitochondrial functions, such as ROS generation, mitochondria to nucleus signaling, mitophagy, and the tethering of signaling molecules to the mitochondrial membrane, have prompted a reevaluation of the mitochondria’s role as organelles involved in modulating signal transduction (Liu and Butow, 2006; Chandel, 2014; Wang and Chen, 2015; Wrobel et al., 2015; Weidberg and Amon, 2018; Innokentev and Kanki, 2021).

In this study, we investigate a novel role of the ETC in glucose derepression. We demonstrate that ETC mutants exposed to glucose fail to grow on galactose, while mutants that have never encountered glucose can utilize galactose as a carbon source. We demonstrate that mutated components of the Snf1 pathway can suppress this phenomenon and show that irregular protein localization within the Snf1 pathway is associated with the reduced ability of ETC mutants to grow on galactose. Collectively, our results reveal a signaling connection by which ETC functionality is critical for appropriate regulation of processes involved in glucose limitation through the Snf1 pathway, suggesting a potentially novel signaling role of the ETC. The mammalian equivalent of Snf1, AMPK, has also been implicated as an important modulator of cancer cell proliferation and suppression (Sadria et al., 2022). With recent evidence demonstrating the metabolic inflexibility of certain cancer cells during glucose to galactose transitions, the results from this study can be used to explore and potentially exploit this phenomenon as a metabolic treatment for cancer cell proliferation (Zheng et al., 2020).

## RESULTS

### Describing the FGD phenotype

It is well documented that respiratory incompetent *Saccharomyces cerevisiae* cells display a low or negligible galactose utilization phenotype. We confirmed that cells treated with ethidium bromide (EtBr) and lacking mitochondrial DNA (mtDNA), known as rho^0^ mutant cells, exhibited a reduced ability to grow on galactose-containing minimal medium after being cultured overnight in glucose (Supplemental Figure 1). Cells cultured on galactose can perform either fermentation or respiration using this carbon source, although it is less preferred as a fermentative substrate compared to glucose (Fendt and Sauer, 2010). While rho^0^ cells cannot respire galactose, they can still oxidize galactose to produce ATP through glycolysis and fermentation. To discern whether the reduced ability of rho^0^ strains to grow on galactose stems solely from respiratory incompetence or is influenced by other cellular networks, we performed a modified experiment: rho^0^ cells were generated while culturing cells exclusively on either glucose or galactose (the mutant cells generated in galactose have therefore never experienced glucose repression). The rho^0^ cells generated after culturing on glucose or galactose were termed rho^0^_glu_ or rho^0^_gal_, respectively. We also included a control where rho^0^_gal_ cells were grown overnight in galactose (glucose derepressing condition) to qualitatively determine growth when rho^0^ cells are not glucose repressed. Both rho^0^_glu_ and rho^0^_gal_ failed to grow when cultured overnight in glucose and plated onto minimal galactose medium. However, rho^0^_gal_ cells, which were exclusively cultured only on galactose and grown overnight in galactose, exhibited full capability to utilize galactose as a carbon source (Supplemental Figure 1). Therefore, the reduced ability of rho^0^ strains to grow on galactose was dependent on whether the cells had previously encountered glucose.

Observed phenotypes of rho^0^ cells could be the result of multiple independent defects such as lowered mitochondrial membrane potential affecting protein import, higher mutation rate, changes in mitochondrial morphology, or alterations in mitochondrial dependent metabolism (Dimitrov et al., 2009; Vowinckel et al., 2021). As such, a number of different potential mechanisms could explain the reduced ability of rho^0^ strains to grow on galactose. To determine whether failure to grow on galactose is specifically a consequence of respiratory incompetence, we evaluated single gene deletions that render the cell respiratory incompetent without affecting as many mitochondrial processes as observed in rho^0^ cells (Lewis et al., 2021). We targeted two nuclear-encoded ETC genes, *COQ2* (subunit within Coenzyme Q) and *COX4* (subunit within cytochrome *c* oxidase, also called Complex IV), to determine whether these ETC mutants display a similar reduced ability to grow on galactose as observed in rho^0^ strains. Heterozygous versions of *COQ2/coq2*Δ and *COX4/cox4*Δ were maintained only on galactose then sporulated and tetrad dissected strictly on galactose-containing media. We refer to the resulting haploid strains as *coq2Δ*_gal_ and *cox4Δ*_gal_ to indicate strains which have been constructed and cultured only in galactose. Importantly, these mutant cells have never encountered environmental glucose (or glucose repression). Subsequent patching, streaking or maintenance of these strains was performed only in galactose-containing media (Figure 1A). Following this, haploid strains were grown overnight either in glucose- or galactose-containing media. Cells were then pronged onto minimal medium containing either glucose or galactose as well as rich medium with glycerol/ethanol to confirm respiratory incompetence of ETC mutants. As observed in Figure 1B, *cox4Δ*_gal_ was incapable of growing on galactose minimal medium when cultured overnight in glucose but could utilize galactose as a carbon source when maintained and precultured in galactose. While wild type cells can switch between conditions of glucose repression and glucose derepression (based on the carbon source present), respiratory incompetent cells fail in this regard. *cox4Δ*_gal_ can efficiently use galactose as a carbon source only when cells have not encountered glucose. Upon glucose repression, *cox4Δ*_gal_ largely lose the ability to grow on galactose. We also observed this phenomenon in a different ETC mutant – *coq2Δ*_gal_ (Supplemental Figure 2), highlighting the general role of ETC dysfunction on glucose derepression rather than subunit-specific roles. We term this phenotype <u>F</u>ailure of <u>G</u>lucose <u>D</u>erepression (FGD). Due to the ease of working with nuclear gene deletion ETC mutants compared to rho^0^ mutants, follow-up experiments were performed with the nuclear gene deletions.

**Figure 1.**
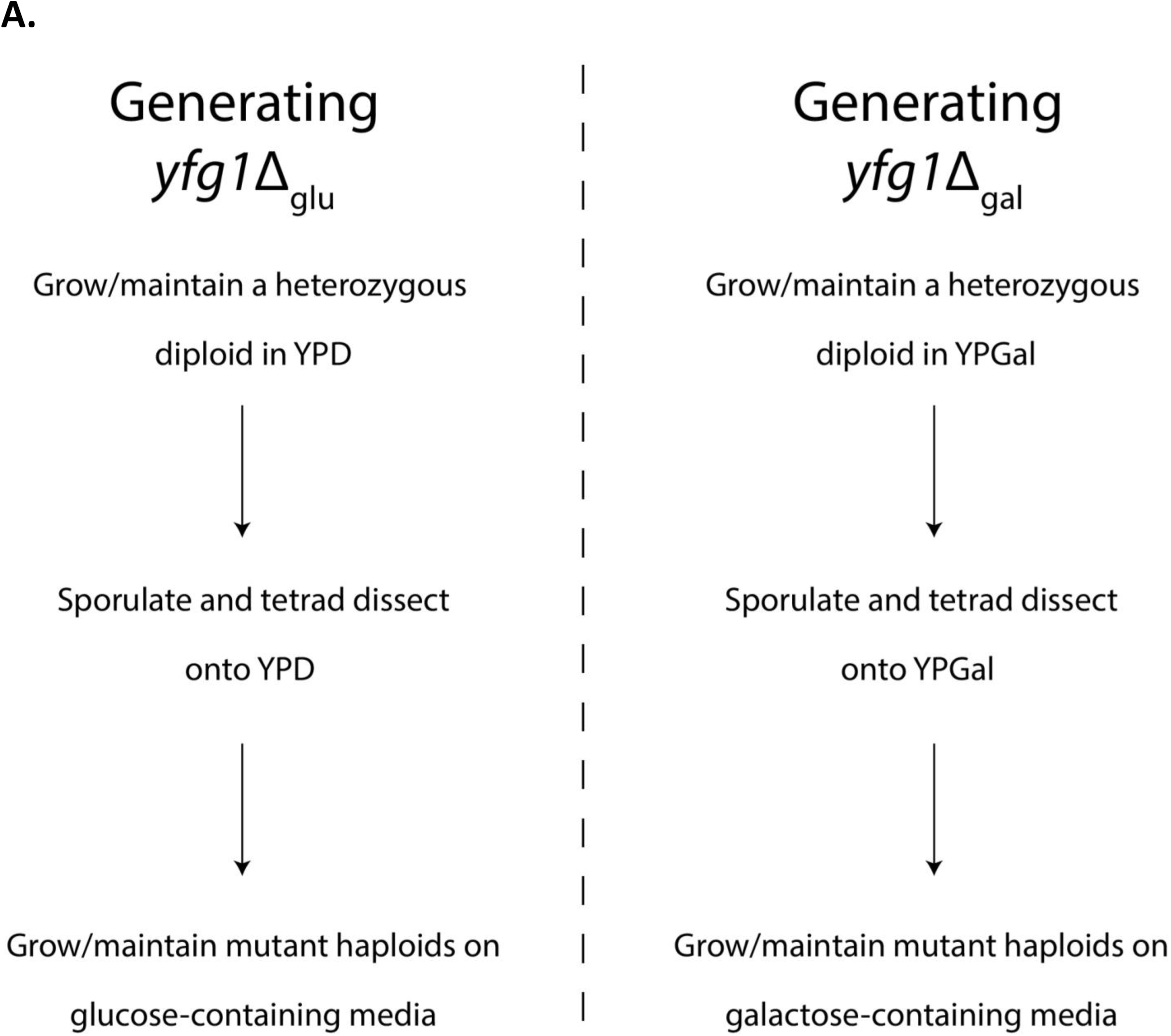
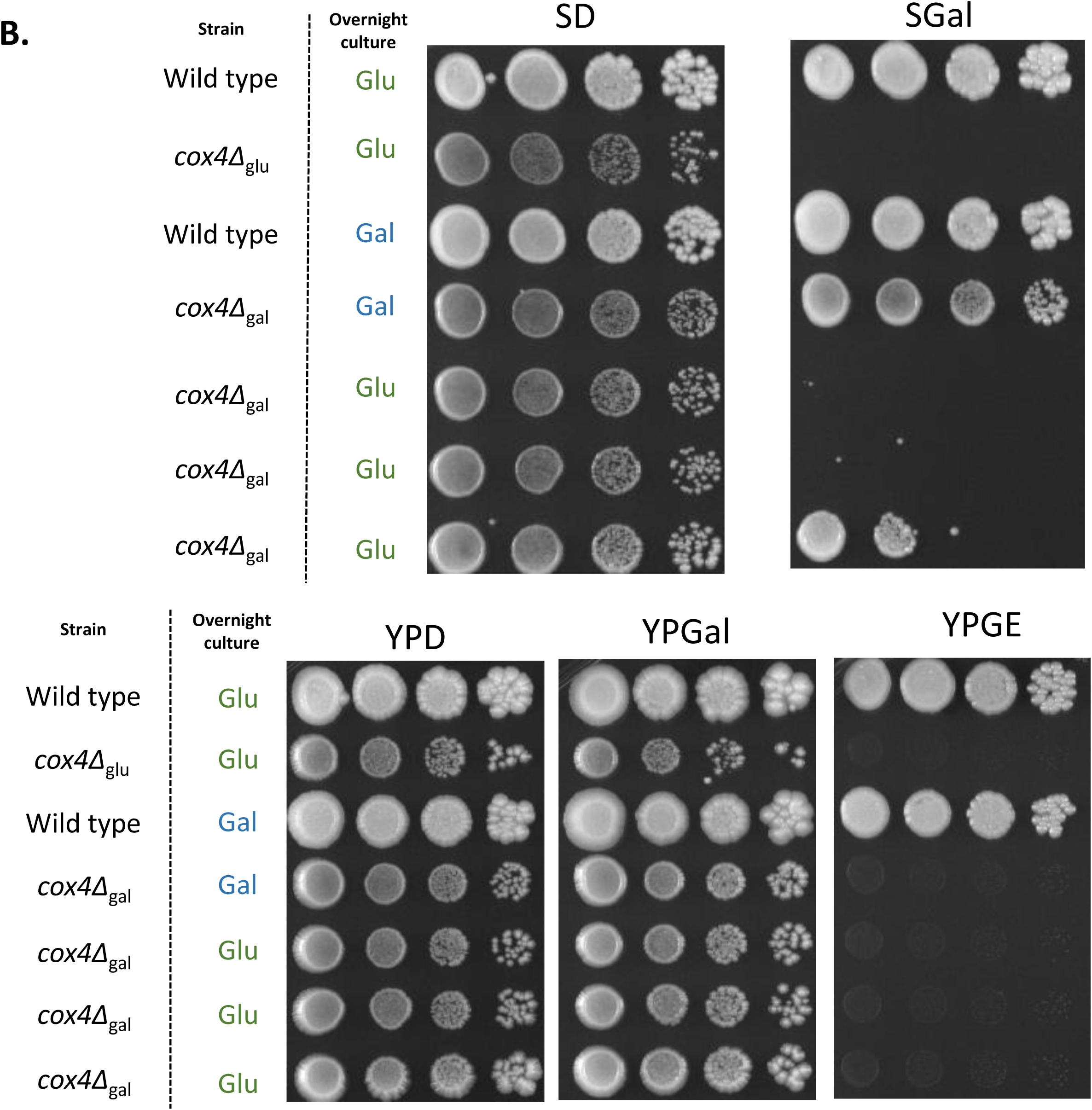
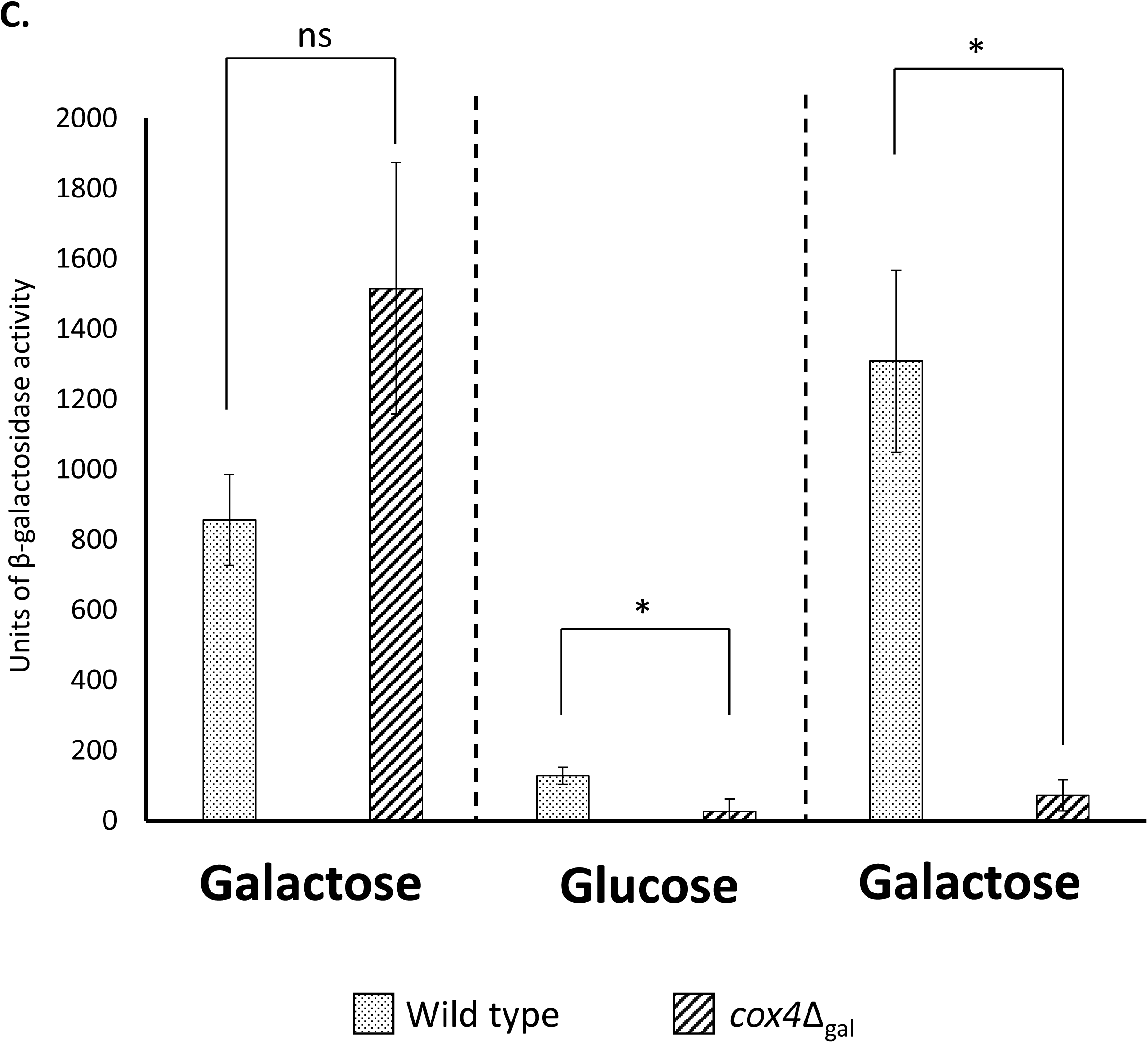
Describing the FGD phenomenon. **(A)** Schematic of strain generation and propagation. Heterozygous diploid strains of the gene of interest (*YFG1*/*yfg1*Δ) were streaked onto either YPD or YPGal. Subsequent strain passages, sporulation and tetrad dissection were strictly performed on the same carbon source. **(B)** Wild type (DBY12000), *cox4Δ* maintained only on glucose (denoted as *cox4Δ*_glu_), *cox4Δ* maintained only on galactose (denoted as *cox4Δ*_gal_) were grown overnight in either SD or SGal as indicated in green and blue text respectively. After overnight growth, 10-fold dilutions were prepared and pronged onto SD, SGal, YPD, YPGal and YPGE. Plates were allowed to incubate at 30°C for 6 days before photographing. **(C)** *ura3*Δ_gal_ and *cox4Δura3*Δ_gal_ transformed with pCM64-*GAL1_pr_-lacZ* were grown overnight in SGal. Cultures were then subcultured into fresh SD. Cultures were allowed to grow for 24h at 30°C before a final subculture into fresh SGal for 24h. Cells were collected from each media change and cell pellets were stored at -80°C until β-galactosidase activity was assessed. Error bars denote standard deviations with 3 biological replicates. Asterisks indicate significant statistical difference (*p* < 0.05; Welch two-sample t-test) between wild type and *cox4*Δ_gal_. ns = not significant (p ≥ 0.05).

### Variability and quantification of the FGD phenotype

We observed a degree of variability in the FGD phenotype based on differing levels of galactose growth among replicates. Out of the three biological replicates of *cox4Δ*_gal_ cultured overnight in glucose shown in Figure 1B, two replicates largely exhibited an inability to consume galactose whereas a small fraction of cells in the third replicate were able grow. Additionally, we also observed that FGD occurred only when strains were pronged on minimal medium containing galactose; ETC mutants’ failure to grow on galactose was not evident when cells were pronged on rich medium (Figure 1B – bottom panel). All *cox4Δ*_gal_ and *coq2Δ*_gal_ replicates grown overnight in glucose were able to grow on rich medium containing galactose. Previous studies have also reported phenotypic variability in growth ability between rich and minimal media (Gibney et al., 2020; Rodríguez-López et al., 2023). In contrast to minimal medium, rich medium is considered chemically undefined due to the yeast extract and bacto peptone components. Factors including nitrogen source (ammonia vs. amino acids) and pH also differentiate rich media from minimal media. To understand what component of rich growth media enables glucose repressed ETC mutants to grow on galactose, we individually tested each ingredient when added to SGal. Surprisingly, only the addition of yeast extract to SGal was able to reverse the FGD phenotype in ETC mutants (Supplemental Figure 3). Addition of bacto peptone (a type of proteinaceous lysate of animal origin) or a mixture of amino acids (the SC supplement mix) were insufficient for ETC mutants to overcome FGD. Moreover, altering the pH of YPGal (pH∼6.5) to that of SGal (pH∼5.5) did not cause FGD in rich medium (Supplemental Figure 3).

As the observation of FGD relies on galactose metabolism, we used a *GAL1-lacZ* reporter to quantify galactose-regulated gene expression between ETC mutants and respiratory competent wild type cells in FGD conditions. Both wild type and *cox4Δ*_gal_ were initially able to grow on galactose and exhibited comparable β-galactosidase levels (Figure 1C). When these strains were transferred to minimal glucose medium, both strains exhibited low levels of β-galactosidase activity as expected as a result of glucose repression, although wild type still had significantly higher residual β-galactosidase activity compared to *cox4Δ*_gal_. After growth on glucose, we transferred the strains back to galactose. As seen in Figure 1C, β-galactosidase activity of wild type rebounded to levels typical of initial growth in galactose while activity of *cox4Δ*_gal_ remained negligible. This result indicates that ETC mutants failure to grow in minimal galactose medium is not simply a metabolic failure. The observed gene expression defects suggest signaling dysregulation, and that components of the electron transport chain, or perhaps intermediate metabolites produced, have a role in relieving glucose repression.

### The FGD phenotype is not strain-specific

The impact of strain background on phenotypic response can impede our ability to characterize a genetic variant (Galardini et al., 2019). Strain-specific phenotypes in glucose metabolism have already been documented – for example, variable growth of a trehalose-6-phosphate synthase mutant (*tps1*Δ) on glucose and fructose was caused due to an S288C-specific persister-like state of *tps1*Δ which relied on *MKT1* function (Gibney et al., 2020). These observations indicate value in evaluating a novel phenotype to determine whether it is strain-specific or more generally conserved. As all the experiments described thus far were conducted in a laboratory *GAL+ HAP1*+ S288C-derivative strain, we sought to determine whether FGD was an S288C-specific phenotype. To assess this, we expanded our yeast panel to include a subset of wine yeast isolates as well as other commonly used strains. In addition to S288C, we included two commercial wine yeast isolates (Simi White and CSM), an oak tree isolate (YPS1000), a vineyard isolate (Bb32) and a different laboratory strain, W303. rho^0^_gal_ versions of all strain backgrounds were prepared by culturing and maintaining strains strictly on galactose and treating cells with ethidium bromide as described in Materials and Methods. Strains with and without intact mtDNA were then grown overnight in either glucose (to initiate glucose repression) or galactose and then pronged onto SD and SGal. We observed that all strains, irrespective of strain background, displayed FGD as demonstrated by reduced ability to grow on SGal after being glucose repressed (Figure 2). However, galactose-maintained strains which were never glucose repressed (overnight growth in galactose) were able to consume galactose. These results suggest FGD is a conserved phenotype to some degree, rather than specific to the S288C genetic background.

**Figure 2.**
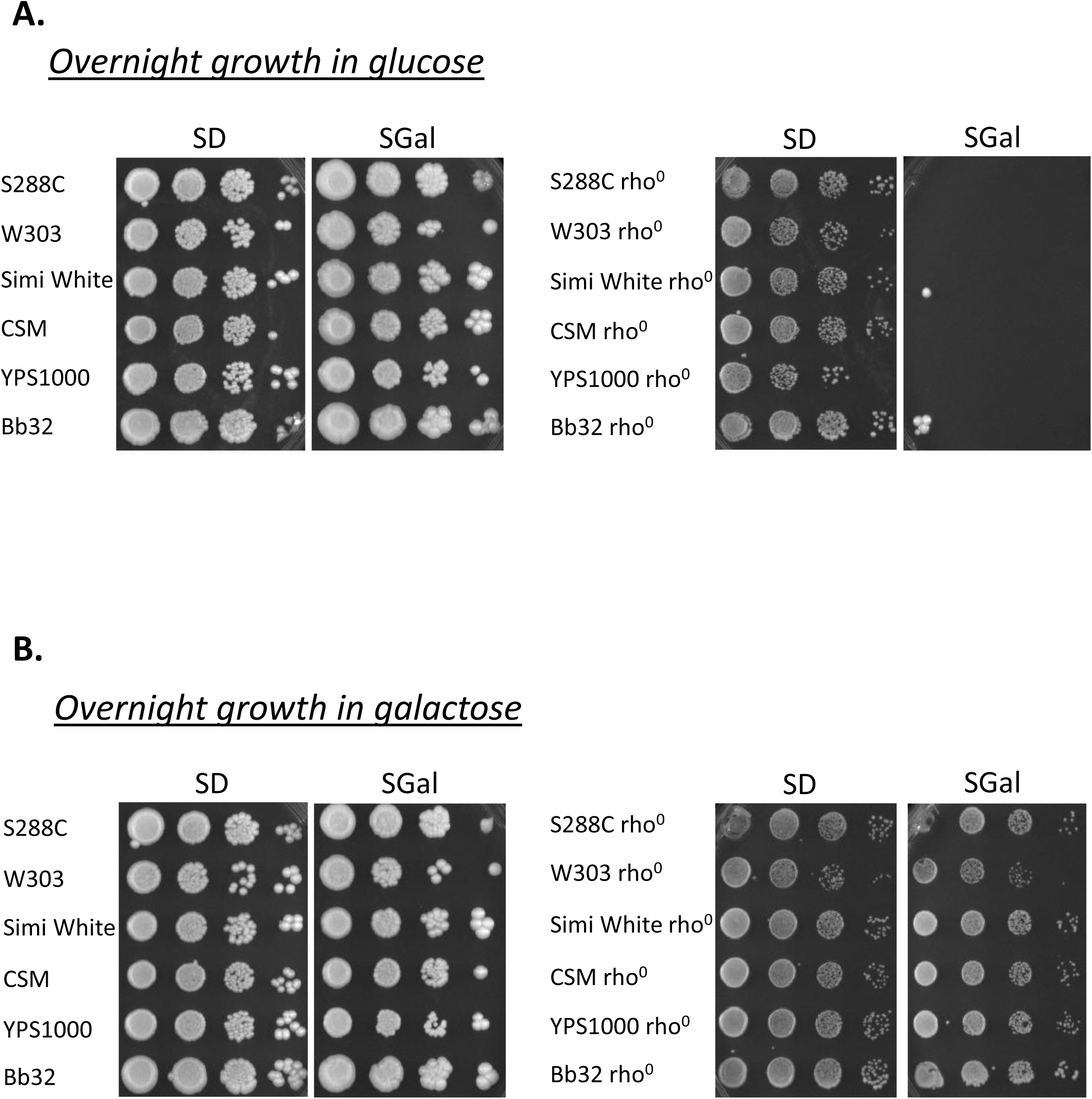
FGD phenotype observed in wild yeast isolates. S288C-derivative DBY12007, W303, Simi White, CSM, YPS1000 and Bb32 strains were treated with EtBr to generate rho^0^ respiratory incompetent cells while ensuring strains were cultured and maintained only on galactose containing growth media. Respiratory competent and incompetent (rho^0^) versions of the strains, maintained only on galactose, were grown overnight in either minimal glucose (**A**) or galactose (**B**) and 10-fold serial dilutions were pronged the next day onto SD and SGal. Plates were incubated at 30° and photographed after 6 days of incubation. Shown are representative images of at least 3 biological replicates.

### Loss of ETC function in glucose repressed cells causes dysregulation of the Snf1/AMPK pathway

Taken together, characterization of the FGD phenotype suggested that glucose sensing/signaling is defective in ETC mutants in low glucose conditions. We therefore sought to evaluate whether the Snf1 branch of glucose sensing/signaling is functioning properly in ETC mutants. The heterotrimeric Snf1 kinase is the yeast ortholog of mammalian AMP-activated protein kinase (AMPK). While AMPK activity is modulated by declining energy levels and concomitant increased AMP levels, the Snf1 kinase is activated upon glucose depletion by an increase in ADP levels (Hedbacker and Carlson, 2009; Hardie et al., 2011; Mayer et al., 2011; Broach, 2012). The Snf1 complex regulates transcription of a large set of genes including those involved with alternative carbon source utilization (such as galactose and sucrose), gluconeogenesis and respiration related genes in addition to stress response genes (Hedbacker and Carlson, 2009). A key downstream effector of Snf1 kinase is the Cys_2_-His_2_ zinc-finger transcription factor, Mig1. Under high glucose conditions, dephosphorylated Mig1 is retained in the nucleus repressing promoters of glucose repressible genes such as those of sucrose and galactose metabolism via recruitment of the transcriptional corepressor Ssn6-Tup1 complex (Nehlin et al., 1991; Vallier and Carlson, 1994; Treitel and Carlson, 1995; DeVit et al., 1997). Additionally, Hxk2 and Mig1 are reported to form a heterodimer to perform transcriptional repression, with Hxk2 playing a critical role in regulating Mig1 phosphorylation (Moreno and Herrero, 2002; Ahuatzi et al., 2007). Under high glucose conditions, inactive dephosphorylated Snf1 is retained in the cytoplasm. Upon glucose limitation, active phosphorylated Snf1 kinase is translocated to the nucleus and deactivates Mig1 by phosphorylating different serine residues resulting in export of Mig1 to the cytoplasm via the Msn5 exportin (Treitel et al., 1998; DeVit and Johnston, 1999; Smith et al., 1999).

To probe whether Snf1 kinase pathway activity was impaired in ETC mutants during glucose derepression, we first evaluated whether perturbing pathway components that act as negative regulators of glucose derepression suppresses FGD. We constructed individual *MIG1* and *HXK2* deletions in a *coq2Δ*_gal_ and *cox4Δ*_gal_ background, grew the strains overnight in glucose and pronged cells onto glucose- and galactose-containing minimal medium. We observed that galactose maintained ETC mutants containing either *mig1*Δ or *hxk2*Δ deletion were able to grow on galactose-containing medium better than *coq2Δ*_gal_ and *cox4Δ*_gal_, partially suppressing FGD, although growth did not reach wild type levels (Figure 3A and 3B). Moreover, ETC mutants with a *HXK2* deletion showed a weaker FGD suppression as compared to mutants with a *MIG1* deletion. Hxk2 is thought to function as an interacting partner of Mig1 and does not directly repress transcription of glucose repressed genes in contrast to Mig1. Thus, the relative contribution of Hxk2 to FGD may be lower compared to Mig1. These results suggest that FGD in ETC mutants is connected to the Snf1 signaling pathway. Further, these results suggest that galactose growth observed for a minority of cells in FGD plating assays may be due to spontaneous loss-of-function mutations in components of the glucose sensing/signaling network.

**Figure 3.**
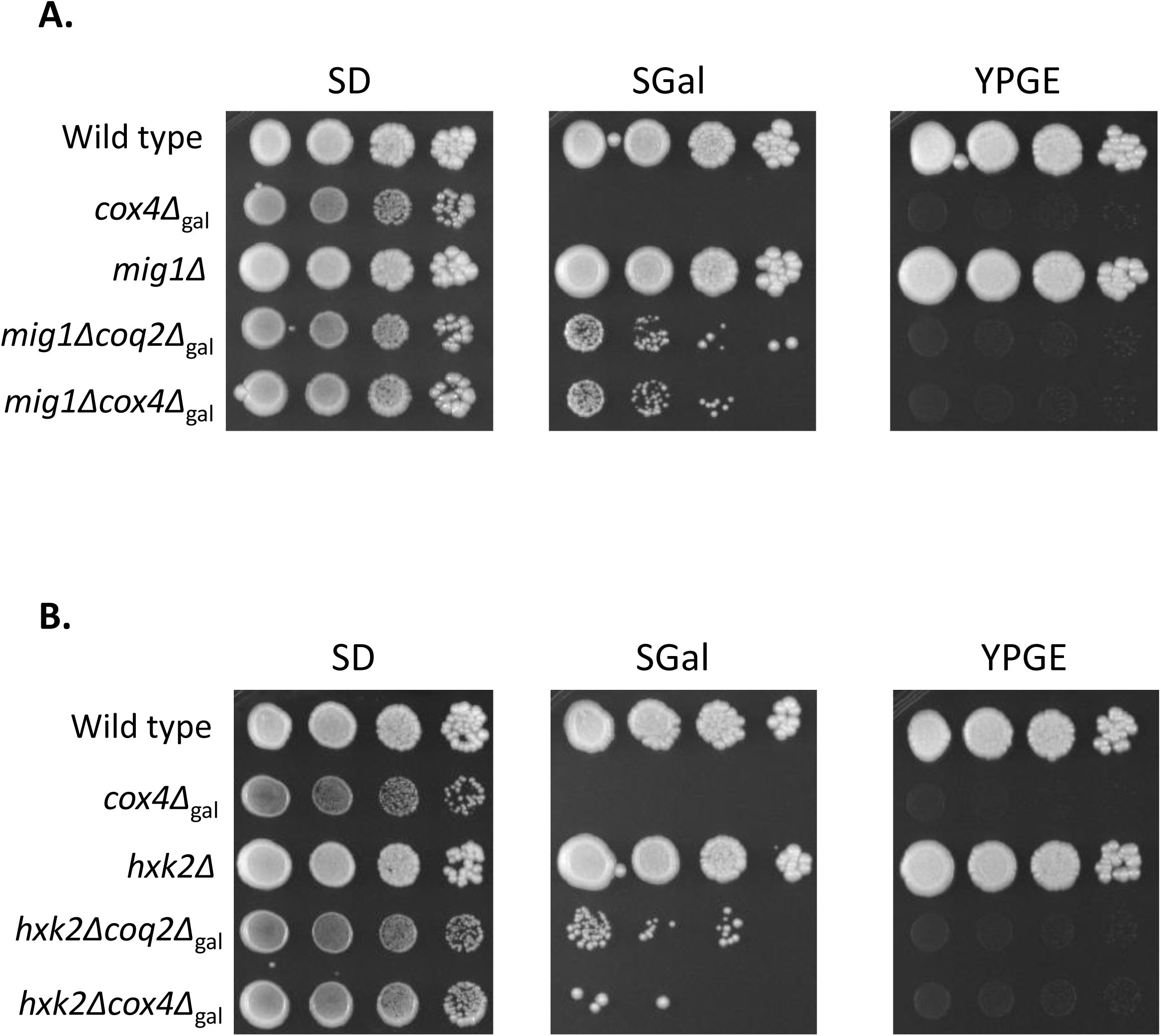
Suppressors of the FGD phenotype include components of the Snf1-Mig1 pathway. **(A)** Wild type (DBY12000), *cox4*Δ_gal_, *mig1*Δ*, mig1*Δ*coq2*Δ_gal_ and *mig1*Δ*cox4*Δ_gal_ were grown overnight in SD and pronged onto SD, SGal and YPGE. Plates were incubated at 30℃ for 6 days before photographing. Shown are representative images of at least three biological replicates. (B) Wild type (DBY12000), *cox4*Δ_gal_, *hxk2*Δ*, hxk2*Δ*coq2Δ*_gal_ and *hxk2*Δ*cox4*Δ_gal_ were grown overnight in SD and pronged onto SD, SGal and YPGE. Plates were incubated at 30℃ for 7 days before photographing. Shown are representative images of at least three biological replicates.

To further evaluate the role of the Snf1 pathway in ETC-mediated FGD, C-terminal fluorescent mNeonGreen protein fusions to Mig1 and Snf1 were used to follow protein localization when cells were cultured under different carbon source conditions. To evaluate protein localization in FGD, all steps of fluorescent strain production, growth and maintenance were performed in galactose-containing media before performing experiments. Additionally, we used an enzyme localized to the nuclear envelope to visualize the nucleus (Hmg1, an HMG-CoA reductase, expressed with a C-terminal fluorescent mRuby2 protein fusion) (Koning *et al*., 1996). To evaluate localization in glucose-repressed conditions followed by derepressed conditions, strains were initially grown in galactose before switching cells to glucose-containing media (SD). After cells reached exponential growth phase in SD, strains were transferred back to galactose and samples were taken at indicated time points. Figures 4A-C demonstrates localization of Mig1 under high glucose conditions (growth in glucose) and when switched to low glucose conditions (growth in galactose) in wild type, *coq2*Δ_gal_ and *cox4*Δ_gal_. For wild type cells, Mig1 is localized to the nucleus under high glucose conditions and relocalizes to the cytoplasm within minutes when galactose is abundant (low glucose conditions) as expected (Figure 4A). However, we observed differences in localization patterns of Mig1 between wild type and ETC mutants. Mig1 in ETC mutants is capable of initially localizing to the nucleus under high glucose conditions, similar to wild type Mig1. However, after glucose repression, ETC mutants are unable to fully export nuclear-localized Mig1 to the cytoplasm (Figures 4B-C). As observed in Figure 4B and 4C, under galactose grown conditions, a portion of Mig1 in both ETC mutants is retained in the nucleus although some Mig1 protein also localizes to the cytoplasm. Figure 4D represents a quantification of the distribution of Mig1 between the nucleus and cytoplasm (data from subsequent biological replicates is presented in Supplemental Figure 4). Quantification of images was performed by calculating the ratio of the pixel intensity of fluorescence in the nucleus to the cytoplasm (ratio_nuc/cyt_) as described in Materials and Methods. A ratio_nuc/cyt_ <1 suggests that the protein is primarily localized to the cytoplasm whereas a ratio_nuc/cyt_ >1 is indicative of nuclear localization. A ratio_nuc/cyt_ of ∼1 suggests a similar distribution of the protein between the nucleus and the cytoplasm. During glucose repression, there is a predominant nuclear localization of Mig1 in wild type and the ETC mutants (ratio_nuc/cyt_ varied from 4-6), while during glucose derepression, the average ratio_nuc/cyt_ for wild type was ∼0.5 demonstrating higher cytoplasmic Mig1 localization whereas the ratio_nuc/cyt_ for the ETC mutants was closer to 1 suggesting comparable distribution of Mig1 between the nucleus and cytoplasm. These results indicate that under low glucose conditions, nuclear Mig1 in respiratory incompetent strains may continue to repress genes for alternative carbon source utilization resulting in the reduced ability of strains to activate GAL gene expression and utilize galactose in minimal medium. We also observed that ETC mutants display a gradual increase in Mig1 foci in the cytoplasm during growth in galactose, although the significance of these foci remains unclear (Supplemental Figure 5).

**Figure 4.**
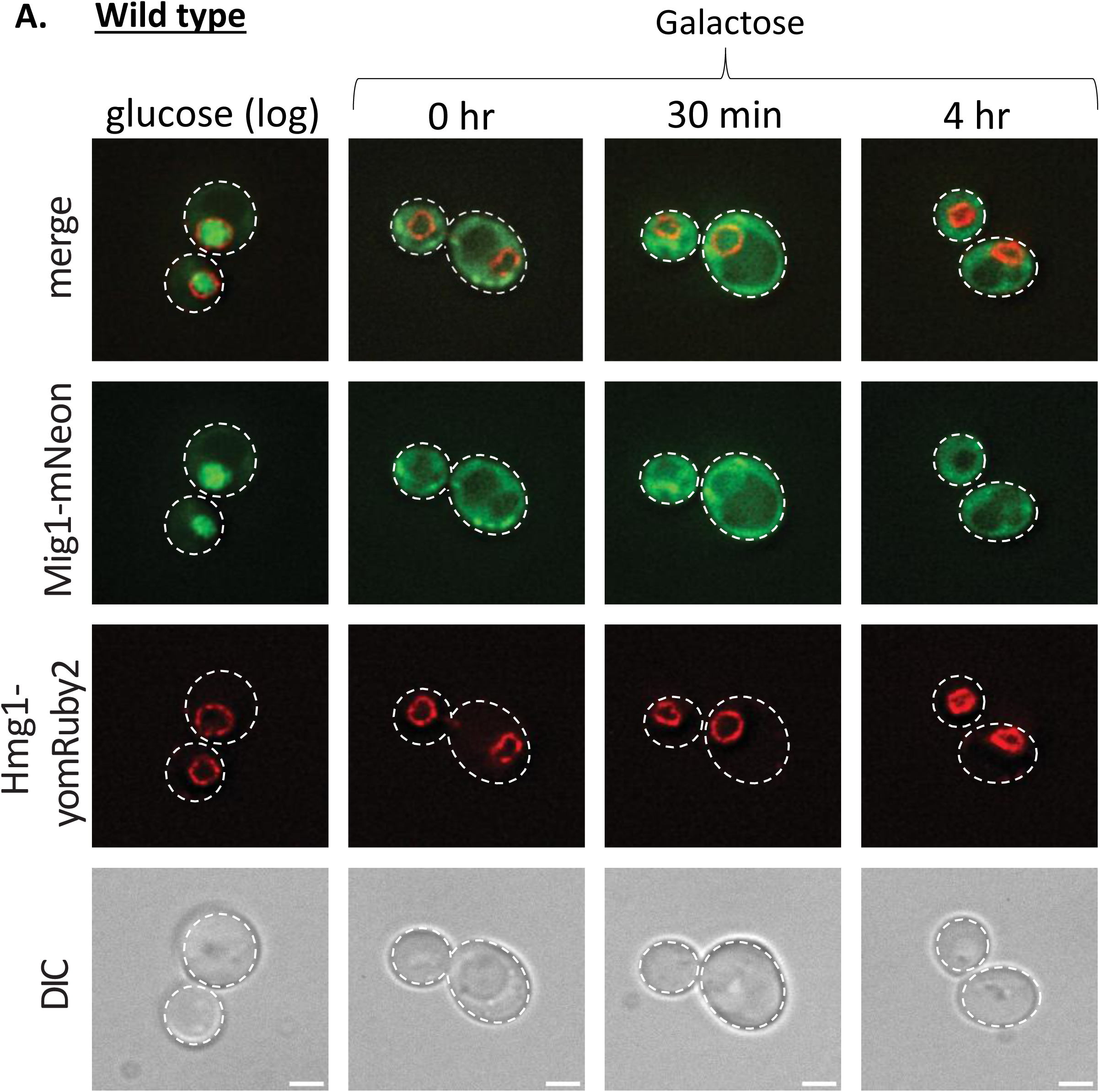
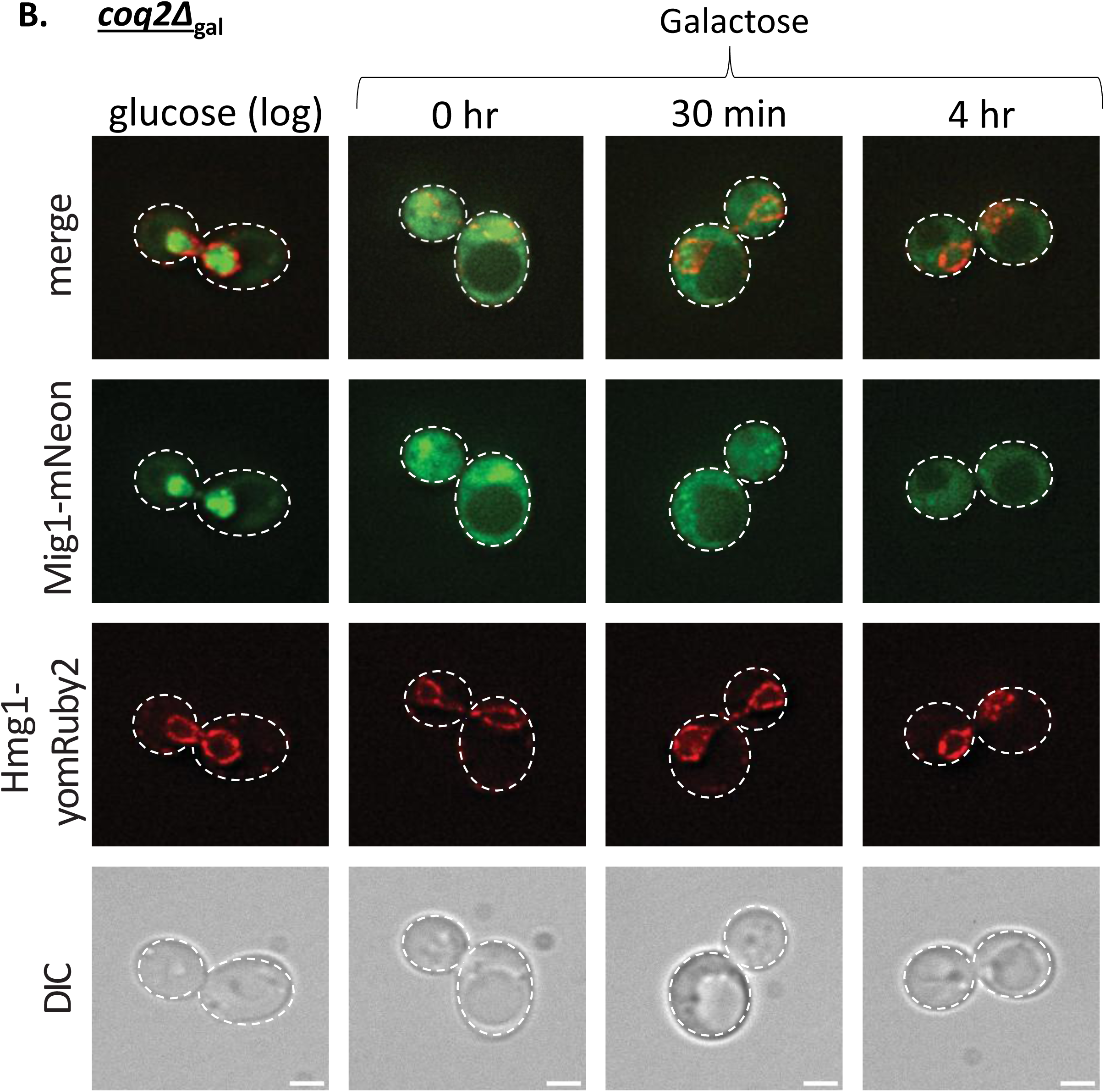
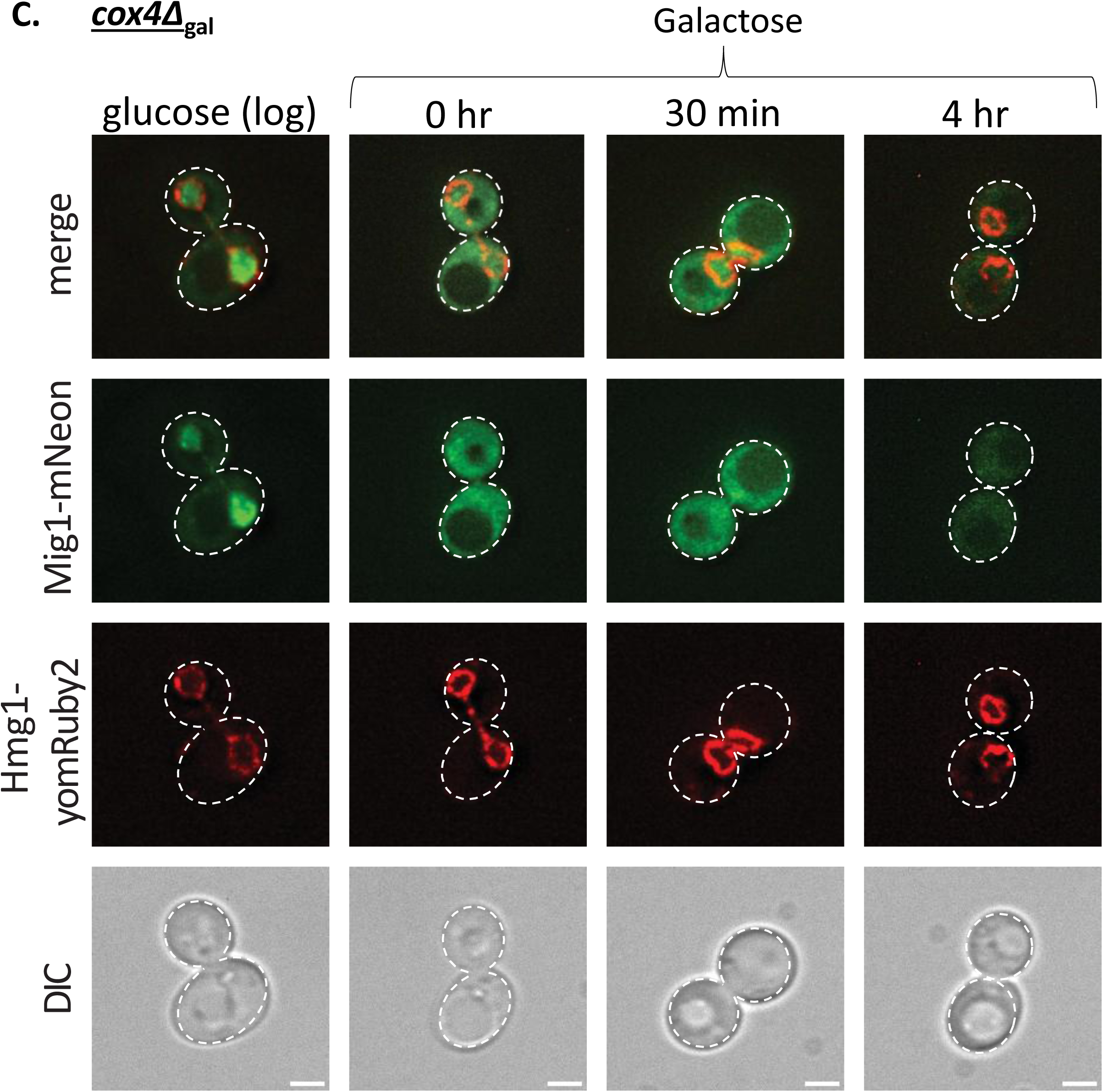
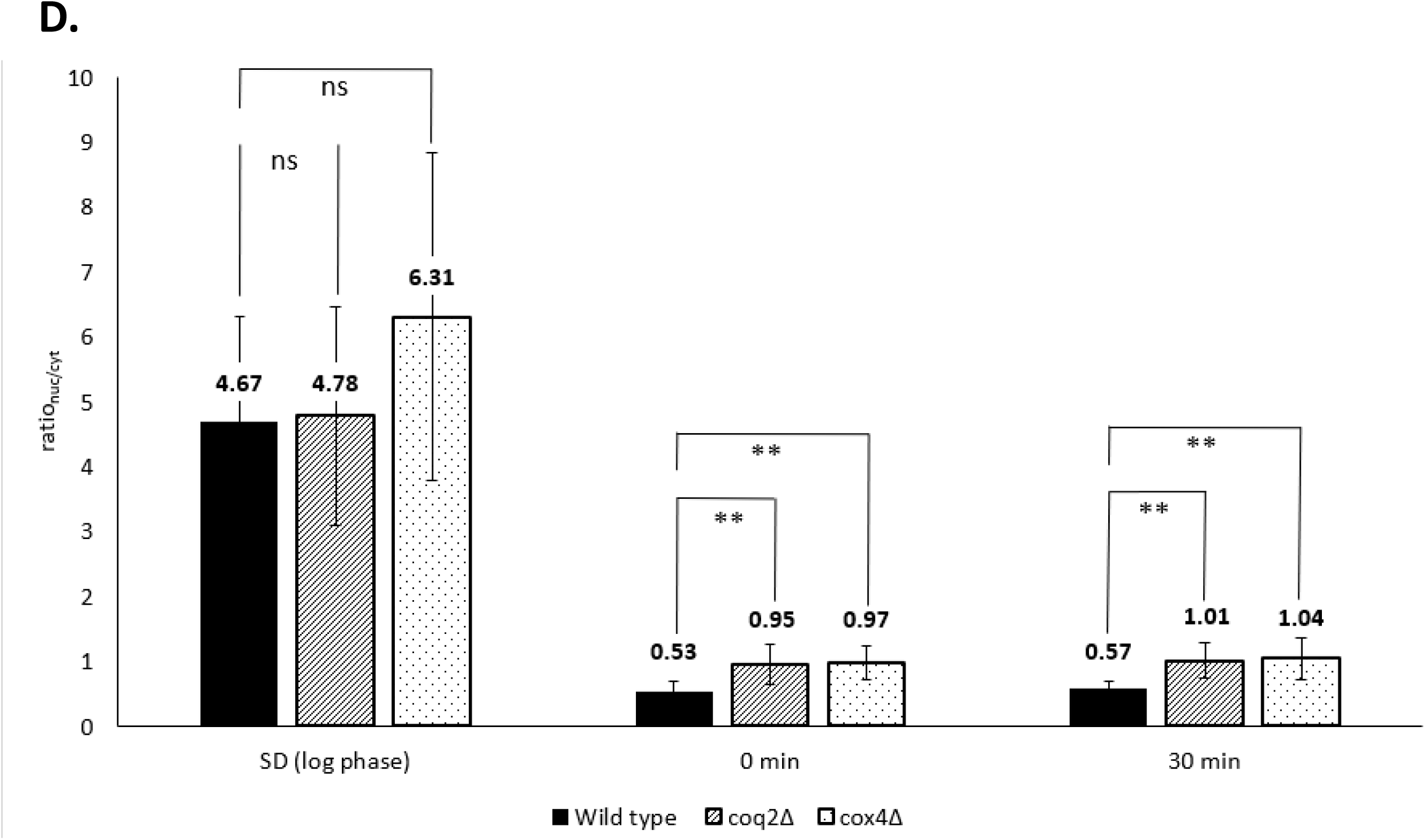
Impaired Mig1 localization observed in ETC mutants during FGD. Localization patterns of Mig1-mNeonGreen in galactose-maintained wild type (**A**), *coq2Δ*_gal_ (**B**) and *cox4Δ*_gal_ (**C**) strains. Cells were grown in SGal overnight before being transferred to SD. Once the culture had reached exponential growth phase in SD, cells were collected by centrifugation, washed with sterile milli-Q water and subcultured into fresh SGalat an initial OD_600_ of 0.2. Images were taken at indicated time points (‘0h’ cells were collected immediately after resuspension into fresh SGal). Three replicates were performed with 20-30 cells visualized and quantified per replicate. Images shown are representative examples from a single replicate. Scale bar = 2µm. (**D**) Distribution of Mig1 protein between the nucleus and cytoplasm was quantified using ImageJ; average ratio_nuc/cyt_ are shown from a single replicate (quantification of subsequent replicates is plotted in Supplemental Figure 4). *: uncorrected *p* value < 0.05, **: uncorrected *p* value < 0.0001, ns: not significant (*p* ≥ 0.05).

Next, we evaluated whether aberrant Snf1 pathway activity in galactose-grown glucose-repressed ETC mutants was also evident upstream of Mig1 functional localization. We used an mNeonGreen protein fusion with Snf1 to investigate whether dysregulation takes place at the level of the Snf1 kinase complex. Previous fluorescent protein localization studies have shown that the Snf1 kinase localizes to the cytoplasm under high glucose conditions but enters the nucleus under low glucose conditions (reciprocal of Mig1 localization patterns), thus serving as a read-out for Snf1 function (Vincent et al., 2001). Similar to previous localization experiments, we transferred galactose-maintained wild type and ETC mutants between different carbon sources. Under high glucose conditions, Snf1 is diffused within the cytoplasm whereas under low glucose conditions, Snf1 localization to the nucleus increases. For glucose-grown wild type and ETC mutants, Snf1 is retained in the cytoplasm with low fluorescence signal in the nucleus, as expected under high glucose conditions; the ratio_nuc/cyt_ was below one for both wild type and ETC mutants (Figures 5A-D). When these cells were transferred back to galactose, wild type showed a higher accumulation of Snf1 in the nucleus, characteristic of Snf1 localization under low glucose/alternative carbon source conditions, although we also observed a considerable amount of the protein in the cytoplasm (Figure 5A). For glucose-repressed ETC mutants cultured in galactose, Snf1 remained evenly diffused between the cytoplasm and nucleus (ratio_nuc/cyt_ was ∼ 1) rather than accumulating higher levels in the nucleus, as observed in wild type where ratio_nuc/cyt_ > 1 (Figures 5B-D, Supplemental Figure 6). The ability of Snf1 to also localize to the cytoplasm when grown in galactose can be partially explained by the Snf1 kinase complex. Different β subunits direct the kinase complex to different organelles within the cell when grown on non-fermentable carbon sources (Vincent et al., 2001). When grown in ethanol, the Gal83 β-subunit accumulates in the nucleus whereas the Sip1 β-subunit directs Snf1 kinase to the vacuole and the Sip2 β-subunit retains the kinase to the cytoplasm (Vincent et al., 2001). Thus, cytoplasmic localization of Snf1 in galactose-grown wild type is presumably due to association of the complex with either Sip1 or Sip2. The ratio_nuc/cyt_ of wild type during glucose derepression (or galactose growth) was consistently higher than 1.3 suggesting higher nuclear localization while for ETC mutants the ratio_nuc/cyt_ was ∼1 suggesting failure of Snf1 to fully localize to the nucleus (Figure 5D). Taken together, the observed uncharacteristic localization patterns of Mig1 and Snf1 in ETC mutants during glucose derepression suggests that repression of alternative carbon source utilization genes seems to be permanently active, leading to the reduced ability of ETC mutants to utilize galactose when cultured in minimal medium.

**Figure 5.**
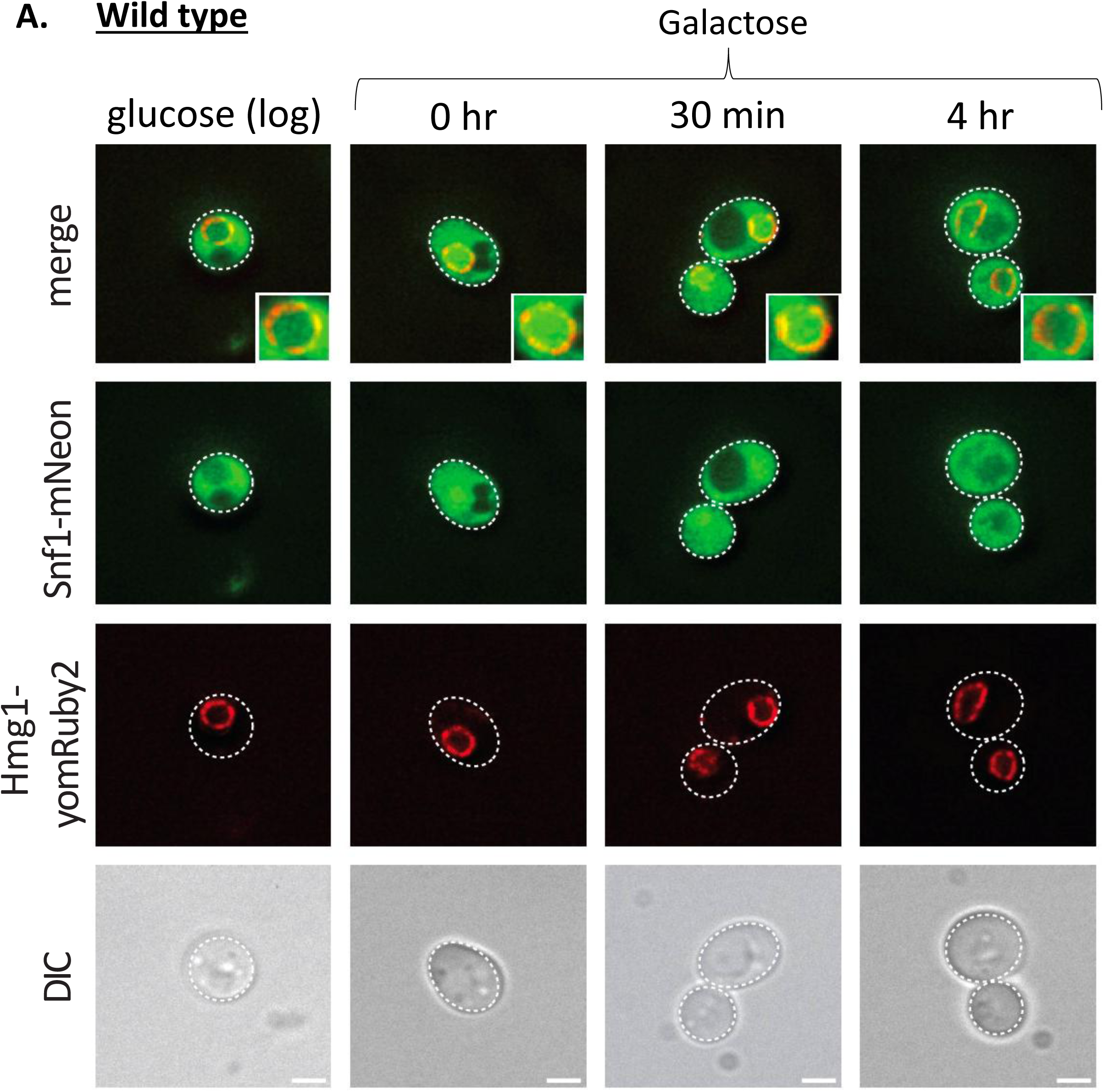
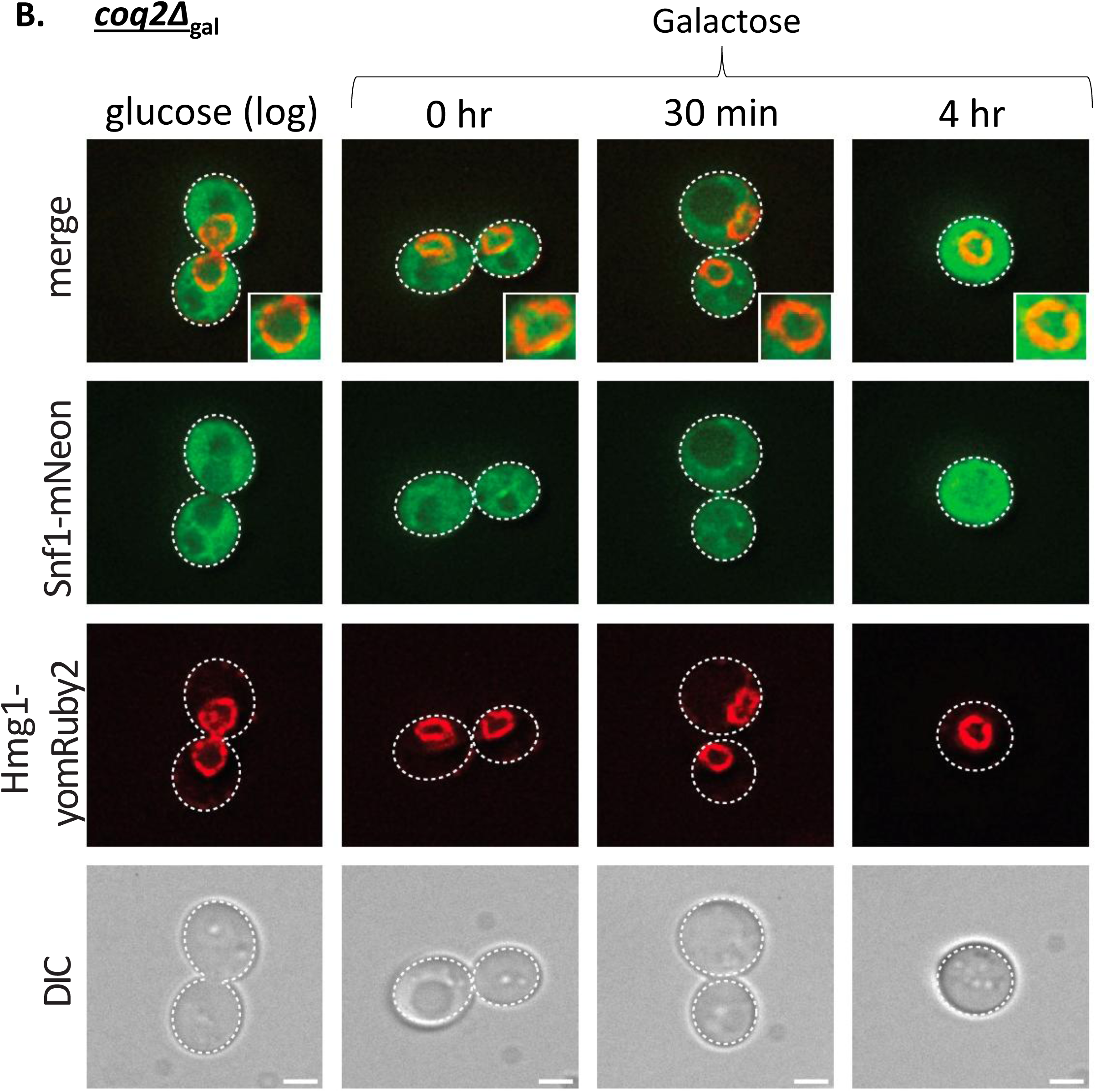
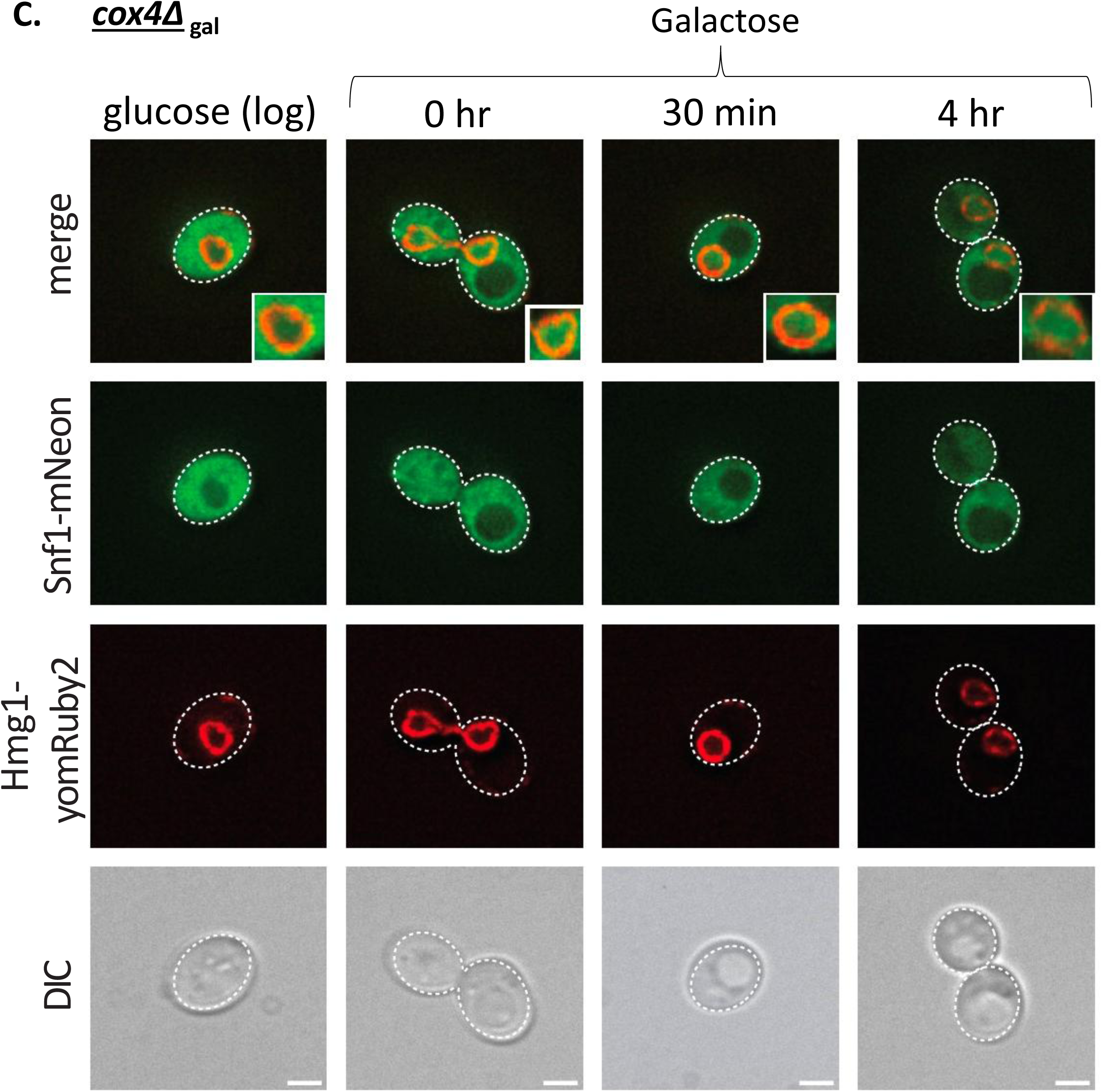
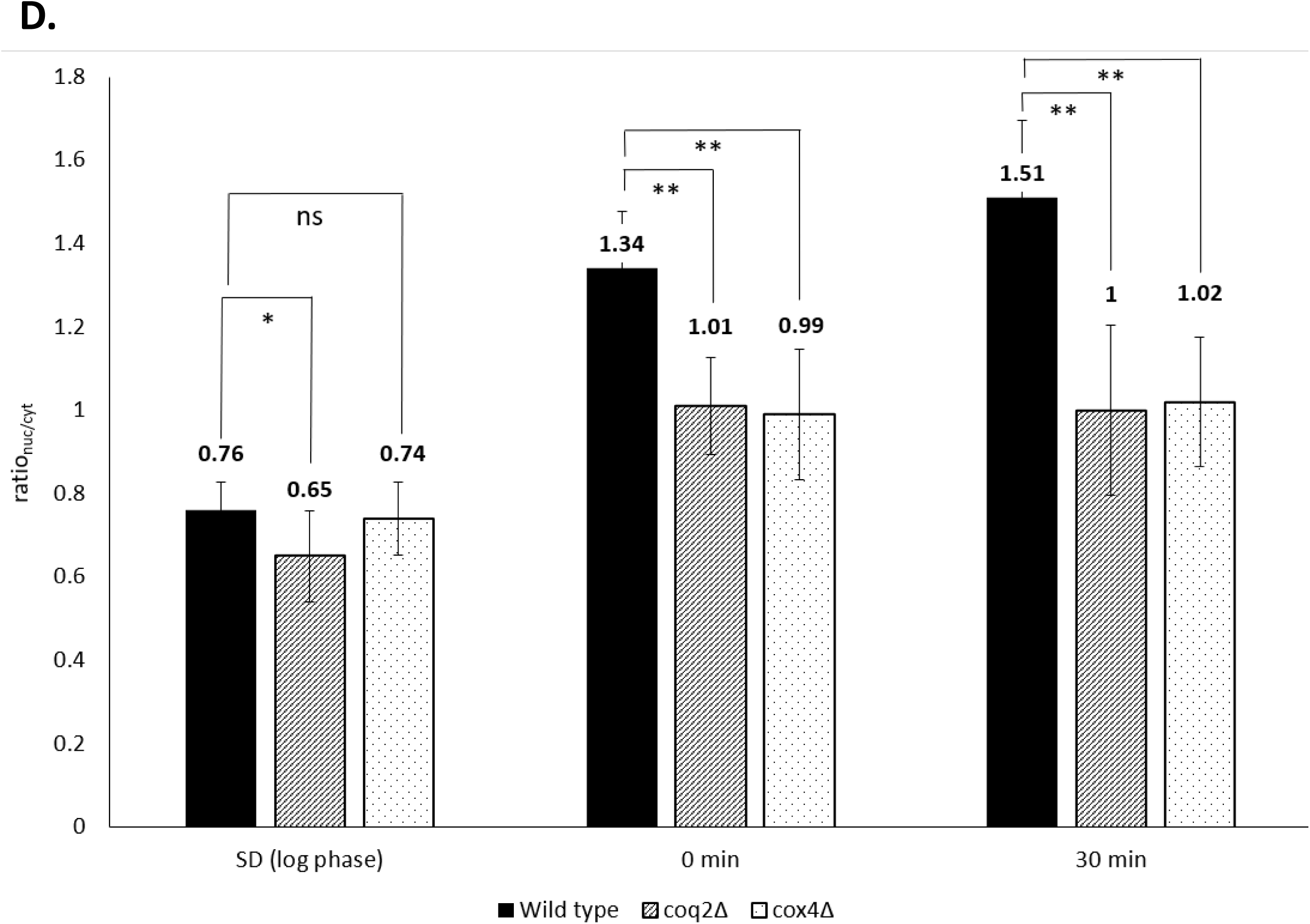
Impaired Snf1 localization observed in ETC mutants during FGD. Localization patterns of Snf1-mNeonGreen in galactose-maintained wild type (**A**), *coq2Δ*gal (**B**) and *cox4Δ*_gal_ (**C**) strains. Cells were grown in SGal overnight before being transferred to SD. Once the culture had reached exponential growth phase in SD, cells were collected by centrifugation, washed with sterile milli-Q water and subcultured into fresh SGalat an initial OD_600_ of 0.2. Images were taken at indicated time points. Three replicates were performed with at least 20-30 cells visualized per replicate. Images were taken at indicated time points (‘0h’ cells were collected immediately after resuspension into fresh SGal). Three replicates were performed with 20-30 cells visualized and quantified per replicate. Images shown are representative examples from a single replicate. Scale bar = 2µm. (**D**) Distribution of Snf1 protein between the nucleus and cytoplasm was quantified using ImageJ; average ratio_nuc/cyt_ are shown from a single replicate (quantification of subsequent replicates is plotted in Supplemental Figure 6). *: uncorrected *p* value < 0.05, **: uncorrected *p* value < 0.0001, ns: not significant (*p* ≥ 0.05).

## DISCUSSION

Here we present a novel regulatory connection for a functional ETC in low glucose conditions through activation of the Snf1 pathway. We confirmed previous observations that respiratory incompetent rho^0^ strains exhibited a reduced ability to utilize galactose as a carbon source. However, these strains are capable of metabolizing galactose when maintained strictly on galactose medium. The occurrence of FGD was not limited to a specific strain background but was rather observed across a diverse range of yeast strains. To evaluate effects of FGD on gene expression, β-galactosidase activity was used to measure *GAL1* expression levels between wild type and ETC mutants. We observed that after glucose repression, *GAL1* induction was significantly higher in wild type cells compared to ETC mutants when grown in galactose. We probed Snf1 pathway activity by studying localization patterns of Mig1, a key transcription factor directly regulated by the Snf1 protein kinase, and Snf1. Intriguingly, glucose-repressed ETC mutants cultivated in galactose displayed impaired clearance of Mig1 from the nucleus. Collectively, these findings provide evidence for the involvement of the ETC in glucose sensing and signaling networks, particularly through the Snf1 pathway during glucose derepression.

### Glucose-repressed ETC mutants display reduced ability to metabolize galactose

Higher concentrations and/or longer treatment with EtBr results in respiratory incompetent cells with different degrees of mtDNA mutation or complete removal of mtDNA (Leibowitz, 1971). Additionally, while there have been theories by which EtBr selectively targets mtDNA without affecting nuclear DNA, a mechanism has never been empirically determined (Goldring et al., 1970). As our initial observations were with EtBr treated rho^0^ cells, we also constructed specific gene deletions of components in the electron transport chain causing the cell to become respiratory incompetent. This ensured that phenotypes observed were solely due to loss of electron transport chain function. The limited ability of respiratory incompetent yeast strains to utilize galactose as an energy source has been extensively documented. For instance, certain respiratory incompetent versions of industrial yeasts were incapable of metabolizing galactose unlike their parent respiratory competent strains (Spencer et al., 1983). Furthermore, investigations employing ETC mutants have directly implicated the role of the ETC in galactose utilization. Quarterman *et al*. demonstrated that the *cox9*Δ mutant, respiratory incompetent due to a subunit deletion in cytochrome *c* oxidase (Complex IV), was unable to utilize galactose (Quarterman et al., 2016). Similarly, van den Brink *et al*. reported that the *rip1*Δ mutant, lacking the Rieske Fe-S protein in Complex III, exhibited a reduced ability to consume galactose after growth in a glucose-limited chemostat, aligning with current observations (van den Brink et al., 2009). Our previous research also highlighted the poor growth of ETC mutants on galactose medium after overnight growth in glucose, indicating their reduced capacity to utilize galactose (Lewis et al., 2021). We initially attributed this phenomenon to ETC mutant’s abrogated respiratory capacity. However, the finding that ETC mutants can effectively use galactose when maintained exclusively on galactose suggests that the underlying cause of this functional growth defect lies in dysregulation of glucose sensing and signaling networks rather than galactose metabolism per se. To assess the impact of this defect, we quantified glucose repression/derepression using the β-galactosidase assay, which showed that glucose repressed galactose grown wild type cells exhibited nearly twenty times higher β-galactosidase activity compared to the *cox4Δ*_gal_ mutant, illustrating the effect of FGD on cellular physiology at the transcriptional level of genes associated with alternative carbon source utilization.

### Impaired Snf1/AMPK pathway activity causes FGD in ETC mutants

In light of the well-established role of the Snf1 pathway in the process of glucose derepression and its significance in regulating carbon source utilization, our focus turned towards investigating its potential connection to FGD. To assess this, we employed fluorescent-tagged Mig1-mNeonGreen and tracked its localization in both wild type and ETC mutant strains cultured in galactose after glucose repression. Mig1, a transcription factor, normally localizes within the nucleus under conditions of high glucose concentration, where it actively represses genes involved in alternative carbon source utilization. However, upon glucose depletion or exposure to alternative carbon sources like galactose, Mig1 undergoes phosphorylation, rendering it inactive, and is subsequently exported out of the nucleus, localizing to the cytoplasm (DeVit et al., 1997). Leveraging Mig1-mNeonGreen as an indicator of Snf1 pathway activity, we made several key observations. Initially, all strains exhibited nuclear localization of Mig1 when cultures were grown in glucose. However, upon transfer to galactose, wild type strains demonstrated exclusive cytoplasmic localization of Mig1 (Figure 4A). In contrast, ETC mutant strains exhibited impaired export of Mig1 from the nucleus, resulting in incomplete cytoplasmic localization (Figure 4B-D). Mislocalization of Mig1 indicates that FGD in ETC mutants stems from dysregulation of the Snf1 pathway during glucose derepression. Notably, the phenotypic variability observed in FGD, as depicted in Figure 1B, Figure 2, and Supplemental Figure 2 could be explained by incomplete localization of key proteins within the Snf1 pathway – a subset of the glucose-repressed mutant population resumed grow on galactose presumably due to sufficient export of Mig1 from the nucleus.

Following the confirmation of aberrant Mig1 localization in ETC mutants during glucose derepression, we evaluated Snf1 localization as a potential indicator of FGD. In normal circumstances, inactive Snf1 resides in the cytoplasm under high glucose conditions (Vincent et al., 2001). However, upon glucose depletion, active and phosphorylated Snf1 translocates to the nucleus (Vincent et al., 2001). Consistent with this pattern, we observed a pronounced nuclear accumulation of Snf1 in wild type cells cultivated in galactose (Figure 5A). In contrast, ETC mutants exhibit a homogenous distribution of Snf1 between the nucleus and cytoplasm, indicating that the impairment of the Snf1 pathway during glucose derepression occurs at the level of the Snf1 kinase (Figure 5B-D). Nevertheless, we do not disregard the possibility that dysregulation occurs even further upstream, potentially involving Snf1 activation through its upstream kinases (Sak1, Elm1, and Tos3) or negative regulators of Snf1 such as the Reg1-Glc7 protein phosphatase. Moreover, other direct and indirect regulators of Snf1 activity further upstream of Snf1’s directly activating kinases and phosphatases could also be the point within the pathway which communicates with the ETC in low glucose conditions.

### Potential signaling mechanisms between the ETC and the Snf1 pathway under low glucose conditions

Having established that aberrant Snf1 pathway activity appears to underly the phenomenon of FGD in glucose repressed ETC mutants, our next objective was to elucidate the signaling mechanism between the ETC and the Snf1 pathway. We considered whether mitochondrial status, specifically ETC integrity, is communicated to the Snf1 pathway via retrograde signaling in low-glucose conditions. Retrograde signaling is a pathway by which changes to mitochondrial homeostasis are conveyed to the nucleus via multiple transcription factors resulting in upregulation of retrograde responsive nuclear encoded genes (Liu and Butow, 2006). Upon mitochondrial dysfunction, transcription factors associated with the retrograde pathway are activated resulting in increased expression of tricarboxylic acid (TCA) cycle genes and rewiring of nitrogen metabolism (Da Cunha et al., 2015). Because mutants defective in respiration would trigger the retrograde signaling pathway, we asked whether the FGD-illustrated signaling connection between the ETC and the Snf1 pathway took place via the retrograde signaling pathway. Rtg3 is a transcription factor functioning in the retrograde signaling pathway, and together with Rtg1 is responsible for induction of retrograde responsive genes such as *CIT2* (peroxisomal form of citrate synthase) (Jia et al., 1997). Disruption of *RTG3* function abrogates induction of nuclear-encoded retrograde responsive genes and hence pathway activity (Komeili et al., 2000). If the ETC communicates to the Snf1 pathway during glucose derepression via the retrograde signaling pathway, then disrupting retrograde signaling should prevent the signaling just like an ETC mutant and also cause the FGD phenotype. In this manner, a retrograde signaling mutant would not be able to utilize galactose after encountering glucose, similar to the FGD phenotype observed in ETC mutants. We disrupted Rtg3 function by deleting the *RTG3* gene and observed that glucose-grown *rtg3*Δ could subsequently use galactose as a carbon source similar to wild type, in contrast to *cox4*Δ_gal_, and thus did not exhibit the FGD phenotype (Supplemental Figure 7). Another possible signaling mechanism could be related to ADP levels. ADP binding regulates Snf1 activity in a concentration-dependent manner by preventing Thr210 dephosphorylation and Snf1 inactivation (Mayer et al., 2011). As increased ADP levels in low-glucose conditions activate Snf1, we considered the possibility that metabolic dysregulation in ETC mutants could lead to aberrantly low ADP levels in low-glucose, preventing activation of Snf1. Using an LC-MS metabolomics approach, we measured AMP, ADP and ATP levels in wild type and ETC mutants when cultured under conditions with abundant glucose (exponential growth in SD) and with no glucose (incubation in SGal), conditions that mimic those used for fluorescently-tagged Mig1 and Snf1 protein localization. As observed in Supplemental Figure 8, ADP levels were not lower in ETC mutants compared to wild type. ADP levels were higher in ETC mutants in SGal compared to wild type, which would be expected for cells experiencing an energy deficit (AMP levels are also higher, while ATP levels are lower) (Supplemental Figure 8). These results indicate that aberrantly low ADP concentration is not responsible for failed glucose derepression in electron transport chain mutants. In light of this evidence, it can be inferred that other distinct mechanisms are involved in the connection between the ETC and the Snf1 pathway during glucose derepression.

Another potential signaling alternative involves cytosolic pH regulation. Cytosolic pH exerts influence over diverse cellular processes, including metabolism and signal transduction, and is tightly controlled by mitochondrial function, among other regulators (Dechant et al., 2010; Orij et al., 2012). Our previous work demonstrated a correlation between cytosolic pH dysregulation and the shortened chronological lifespan of ETC mutants in minimal medium (Lewis et al., 2021). In line with this evidence, a recent study demonstrated that activity of Snf1 is regulated by a poly-histidine tract present within the pre-kinase domain of the Snf1 protein (Simpson-Lavy and Kupiec, 2022). Activity of Pma1, the plasma membrane proton pump, regulates interactions between the N-terminal (which contains the pre-kinase domain) and C-terminal domain of Snf1 via the poly-histidine tract, suggesting that the Snf1 poly-histidine tract acts as a pH sensing module. Taken together, pH-based regulation is a plausible model to explain FGD, though it remains unclear why pH homeostasis is altered in ETC mutants. Other signaling connections are also possible, including mitochondrial quality control mechanisms such as the mitochondrial compromised protein import response triggered by protein import stress, and specific clearance of damaged mitochondria via mitophagic processes (Shpilka and Haynes, 2018; Weidberg and Amon, 2018; Innokentev and Kanki, 2021).

We have previously shown that ETC mutants are unable to survive starvation in glucose minimal medium. Gene expression analysis revealed that transcription factor activity of Cat8 failed to be activated in ETC mutants during post-diauxic shift (Supplemental Figure 9; replotted figure from Lewis *et al*., 2021 (Lewis et al., 2021). Cat8 is a transcriptional activator that activates the Snf1 pathway and controls activation of several glucose repressible genes. Cat8 is activated when glucose is depleted during diauxic shift and cells switch to a respiratory mode of metabolism, i.e., Cat8 is activated during glucose derepression (Randez-Gil et al., 1997; Haurie et al., 2001). Our current findings serve to mechanistically explain the Cat8-mediated gene expression defect observed post-diauxic shift: Cat8 is a Snf1 target, and Snf1 appears to be less active during low glucose conditions in ETC mutants.

Our findings collectively support a proposed model, depicted in Figure 6, which elucidates the cellular events occurring during the transition from a glucose-repressed to glucose-derepressed state. This transition is accompanied by the activation of the Snf1 pathway. During Snf1 activation, a functional mitochondrial electron transport chain plays a critical role in facilitating Snf1 kinase activity. The failure to achieve correct and complete localization results in the reduced ability of cells to grow on alternative carbon sources, such as galactose. Future work to evaluate this model could focus on independent evaluation of a connection between ETC function and Snf1, perhaps through phospho-site-specific protein analysis of Snf1 from nuclear or cytoplasmic fractions of cells grown in different carbon sources. Further work is also required to precisely determine the molecular mechanism underlying the connection between the ETC and Snf1.

**Figure 6.**
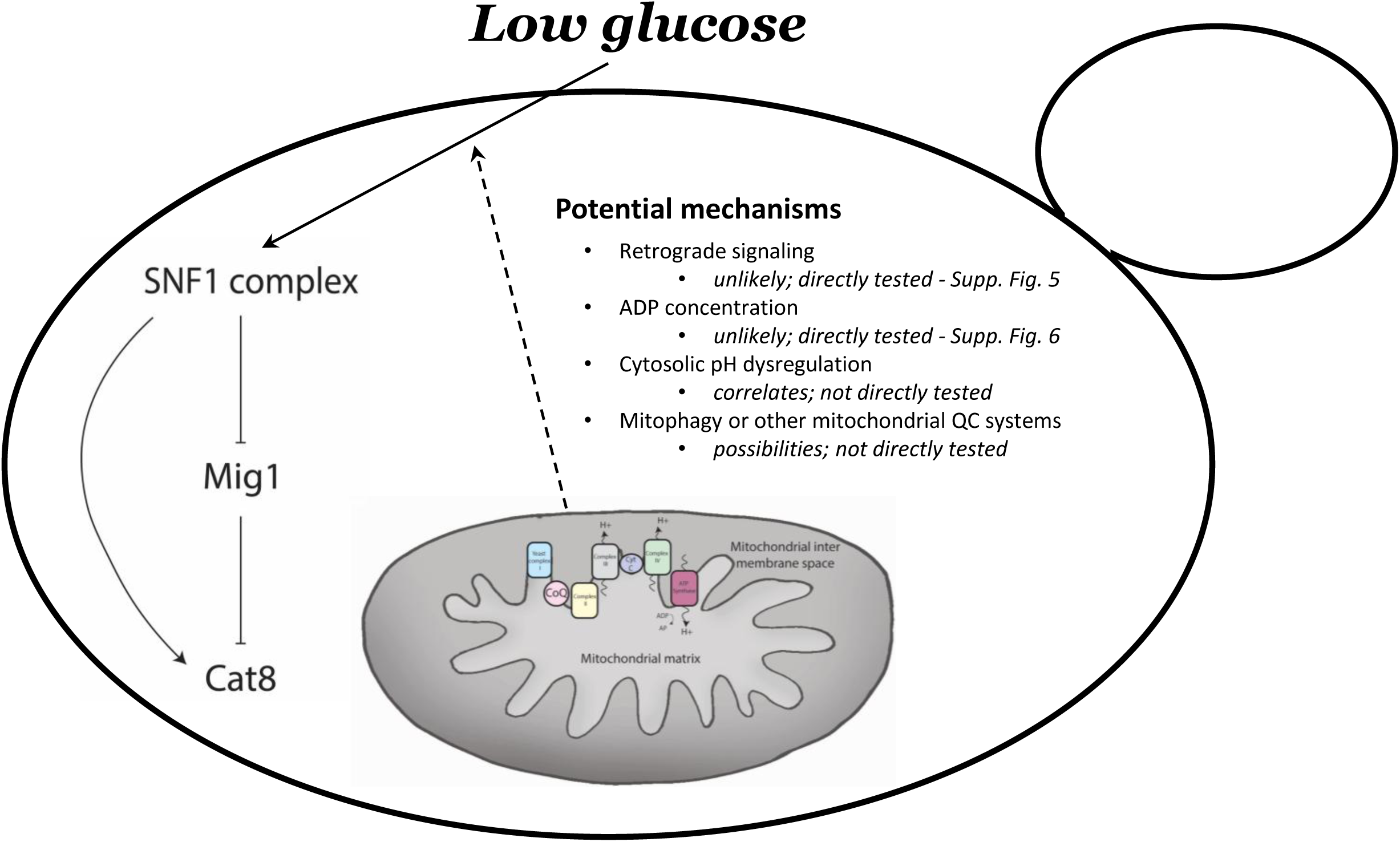
Proposed model for cross-talk between the ETC and the Snf1/AMPK pathway. From the evidence generated, a model for how the ETC regulates the Snf1 pathway under conditions of low glucose is proposed. Components of the ETC, excluding ADP levels or retrograde signaling, are proposed to participate in correct localization of Mig1 to the nucleus under low glucose conditions. Failure to do so results in a dysfunction of alternative carbon source utilization, a phenomenon we term failure of glucose derepression (FGD).

### Potential FGD conservation

AMPK, the mammalian counterpart of the Snf1 kinase, serves as a crucial regulator of energy homeostasis. Similar to the Snf1 complex, AMPK possesses a heterotrimeric structure and is regulated by ADP/ATP and AMP/ATP ratios (Oakhill et al., 2011). The involvement of AMPK in metabolic disorders, including cancer, has garnered considerable attention due to its role as a master modulator of energy balance (Jones et al., 2005; Mihaylova and Shaw, 2011). However, its function in cancer is complex, acting as both a suppressor and activator of cancer cell proliferation (Vara-Ciruelos et al., 2019; Sadria et al., 2022). Notably, pharmacological agents such as metformin, a biguanide commonly used in Type 2 diabetes treatment, and 5-aminoimidazole-4-carboxamide riboside (AICAR), an adenosine analogue, have been shown to activate AMPK and are being explored as potential cancer therapeutics (Mihaylova and Shaw, 2011).

Considering the evolutionary similarities between mammalian and yeast cells with regard to both the ETC and Snf1/AMPK, both functionally and structurally, it is possible that the phenomenon of FGD may extend beyond *S. cerevisiae* to mammalian cells. Our results suggest that ETC function is important for the metabolic flexibility necessary for glucose-repressed, fermentative cells to transition into respiratory cells. Many cancer cells, though not all, exhibit a preference for glucose fermentation to lactate rather than glucose catabolism via respiration, despite the presence of oxygen, a phenomenon known as the Warburg effect (analogous to the Crabtree effect in *S. cerevisiae*)(Shiratori et al., 2019). To examine the potential conservation of the FGD phenomenon, future studies could focus on evaluating the metabolic flexibility and viability of fermentative cancer cells with absent or inhibited ETC function during carbon source transitions. Another connection supporting potential conservation of FGD is based on the observation that human mammary epithelial cells constitutively expressing the AGC-type serine/threonine protein kinase oncogene AKT1 undergo cell death upon transition from glucose to galactose (Casamayor et al., 1999)(Zheng et al., 2020). AMPK is an activator of Akt function under diverse stresses, suggesting that aberrant AMPK activity could contribute to cell death during glucose-galactose transition (Han et al., 2018). This observation aligns with our results relating FGD to Snf1 pathway activity during the glucose-galactose transition.

## CONCLUSIONS

The transition in *S. cerevisiae* from glucose-rich to glucose-poor conditions encompasses various physiological changes that facilitate survival under glucose-depleted conditions. Of particular importance is the upregulation of mitochondrial function, which plays a vital role in efficiently utilizing available carbon sources. While the primary function of the ETC is well-established as an ATP-generating system via oxidative phosphorylation, emerging evidence suggests additional roles for the ETC. In this study, we uncover a novel role of the ETC in regulating glucose derepression via the Snf1 pathway. We observe that glucose repressed ETC mutants display a reduced ability to utilize galactose as a carbon source, a phenomenon we term ‘Failure of Glucose Derepression’ (FGD). Further characterization of FGD reveals abnormal localization patterns of Snf1 and Mig1 in ETC mutants cultured in galactose. Interestingly, we find that dysregulated ADP levels and disruption of retrograde signaling are not involved in mediating the communication between the ETC and the Snf1 pathway during glucose derepression, suggesting an alternative undefined mechanism. The resemblance between FGD and the metabolic inflexibility observed in cancer cells during sugar transition suggests potential conservation of this signaling connection and indicates potential metabolic treatment strategies for inhibiting cancer cell proliferation. Collectively, our findings shed light on a novel mechanism operating during low glucose conditions, highlighting the ETC’s potential signaling role within glucose-related signaling networks.

## MATERIALS AND METHODS

### Yeast media, growth, and culturing

Cells were cultured in either minimal (0.67% Yeast Nitrogen Base with ammonium sulfate without amino acids and 2% glucose or 2% galactose – referred to as SD or SGal respectively), or rich media (2% bacto peptone, 1% yeast extract, 2% glucose or 2% galactose – referred to as YPD or YPGal respectively). To test for respiratory incompetence, strains were grown on either SGE or YPGE with either a minimal or rich media base respectively and 3% ethanol and 2% glycerol added as carbon sources. Percentage indications of media components described refer to weight/volume except for glycerol and ethanol which are listed as volume/volume. Where indicated, cells were washed with filter-purified (18.2 MΩ), autoclaved sterile water (Millipore Milli-Q Advantage A10). Optical density at 600nm (OD_600_) was used to estimate cell concentration (Thermo Fisher Scientific Genesys 6 UV-Vis Spectrophotometer). Differences in strain growth were evaluated by pronging cultures onto solid media. This involved growing overnight cultures in indicated media followed by washing, collecting cells by centrifugation, and diluting cultures to an OD_600_ of 1.0. From this initial concentration, a series of 10-fold serial dilutions were performed in a 96-well plate and then spotted onto indicated growth media using a 48-pin or 96-pin replication tool (Sigma-Aldrich).

Respiratory incompetent rho^0^ versions of strains were generated by ethidium bromide (EtBr) treatment as described previously (Fox et al., 1991). Briefly, this involved growing the strain in minimal medium (supplemented with indicated carbon source; either galactose or glucose) and adding 25 μg/ml EtBr to the culture. Following saturation, cells were sub-cultured into fresh minimal medium with indicated carbon source and EtBr added and grown overnight. Cultures were then streaked for single colonies onto either YPD or YPGal. Lack of mitochondrial DNA was confirmed by PCR using three pairs of oligonucleotides which amplified three separate genes in the mitochondrial genome: *Q0010*, *OLI1*, and *COX3*.

### Yeast strain construction

Strains were constructed in a *GAL^+^*, *HAP1*-repaired, prototrophic derivative of S288C. The original strain was received as a kind gift from the Winston lab, where it was initially designated FY2648 (Hickman and Winston, 2007; Hickman et al., 2011). Plasmids containing different drug-resistance or epitope-tagging cassettes, such as pFA6a-kanMX for kanMX, pAC372 for natAC, pAG32 for hphMX, pUG66 for bleMX, pKT127-NeonGreen-natMX, and pKT209-NeonGreen-CaURA3 were used as templates to generate PCR products (Wach et al., 1994; Lorenz et al., 1995; Goldstein and McCusker, 1999; Gueldener et al., 2002). pAC372 was a kind gift from Amy Caudy; the natAC cassette contains a yeast codon optimized nourseothricin-resistance gene, flanked by the *Ashbya gossypii TEF* promoter and 3’ UTR as present in the other MX cassettes. Targeted gene deletions were made by homologous recombination which included designing primers with 40 flanking base pairs identical to the upstream and downstream regions of the target gene. These primer sets were then used to amplify the targeted cassette from the respective plasmids. PCR products generated in this manner were then transformed into a diploid to obtain a heterozygous strain, which was confirmed by PCR, sporulated and then tetrad dissected to isolate MAT<u>a</u> and MATα segregants. Strain construction was performed by mating, sporulating and tetrad dissection for combinatorial gene deletions. Sporulation involved growing cells to log phase in rich media, collecting cells by centrifugation, washing cells in 1% w/v potassium acetate before resuspending cells in 1% w/v potassium acetate. Cultures were incubated at room temperature on a roller wheel for at least 4 days before tetrad dissection. Tetrad dissection and zygote picking was performed using a tetrad dissection scope fitted with a micromanipulator (Zeiss tetrad dissection microscope). Strains used in this study are listed in Supplemental Table 1.

### Assaying β-galactosidase activity

Strains with a *ura3Δ* and *cox4Δura3Δ* background were transformed with a *GAL1_pr_-lacZ* reporter (pCM64-*GAL1*) which is a 2µ plasmid containing *URA3* as a selection marker. pCM64-*GAL1* was a kind gift from Dr. Kevin Morano and constructed by inserting the *GAL1* promoter in front of a truncated *lacZ* reporter present within the pCM64 plasmid, derived from pLG669-Z (Guarente and Ptashne, 1981). Transformed strains were grown in galactose-containing medium overnight, and sub-cultured the following day into fresh SGal medium at an initial OD_600_ of 0.5. Cultures were allowed to grow overnight before cells were collected and sub-cultured into fresh SD medium at an initial OD_600_ of 0.5. Cultures were again allowed to grow overnight before cells were collected and sub-cultured into fresh SGal medium at an initial OD_600_ of 0.5. After overnight growth in SGal, cells were collected again to assay β-galactosidase activity. Whenever cells were switched to a medium with a different carbon source, 0.5 OD_600_ units of cells were collected and stored at -80℃ until assayed for β-galactosidase activity. At least three biological replicates were assayed. Statistical significance was calculated using the Welch two-sample t-test. Data presented are the mean values, with error bars denoting standard deviation.

β-galactosidase assays to quantify the FGD phenomenon was performed as described in Liu *et al*. (Liu et al., 1999). Collected cells were resuspended in 800 µL Z-buffer (60 mM Na_2_HPO_4_, 40 mM NaH_2_PO_4_, 10 mM KCl, 1 mM MgSO_4_, 50 mM β-Mercaptoethanol). Cells were then permeabilized by adding 50 μL chloroform and 50 μL 0.1% SDS. The assay was initiated by adding 4 mg/ml ONPG to the first tube, vortexing and incubating at 30℃ until a pale-yellow color developed. Reactions were terminated by adding 350 μL of 1M Na_2_CO_3_. Samples were centrifuged and OD_420_ of the yellow supernatant was then measured. Units of β-galactosidase activity was calculated as follows:

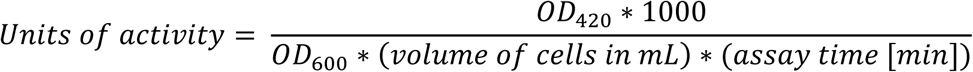

### Fluorescence microscopy

Strains containing fluorescently tagged Snf1 and Mig1 were imaged under different carbon source conditions to study localization patterns of the individual proteins. Snf1 and Mig1 were tagged with fluorescent mNeonGreen whereas Hmg1, an HMG-CoA reductase enzyme involved in sterol synthesis that localizes to the nuclear envelope, was tagged with yeast-codon-optimized mRuby2 for nuclear imaging. Cells were grown in SGal overnight before being sub-cultured in SD medium and incubated until the culture reached an appropriate cell density in exponential growth phase, also called log phase (OD_600_ = 0.2-0.4). Log phase SD cultures were collected by centrifugation, washed, and inoculated into SGal medium at an initial OD_600_ of 0.2. Samples were taken and concentrated at indicated time-points by centrifuging 1 ml of culture for 1 min at 10,000g at 4℃ before cells were resuspended in ∼100 μL of the spent media for live cell imaging.

Imaging was performed on a DeltaVision Elite system (GE Healthcare Life system) fitted with an Olympus IX-71 inverted microscope, a DV Elite complementary metal-oxide semiconductor camera, a 100×/1.4–numerical aperture (NA) oil objective, and a DV Light SSI 7 Color illumination system with Live Cell Speed Option with DV Elite filter sets. Exposure times for the mNeonGreen and mRuby2 channels were 0.5 seconds. Images were captured and deconvolved (conservative setting; seven cycles) using the DeltaVision softWoRx software 7.0.0 (Applied Precision). To quantify protein localization between the nucleus and cytoplasm, deconvolved images were exported to ImageJ. Cells were split into different channels based on the fluorescent marker used. For all images, the brightness for the mRuby2 channel was set to a minimum of 40 and maximum of 2500; for the mNeonGreen channel, the minimum brightness chosen was 100 and maximum was 7000. This allowed for standardization between the different strains, conditions and time points tested. Nuclear and cytoplasmic fluorescence intensity was then measured for each cell using the ImageJ ‘multipoint’ function to select 3 cytoplasmic locations and 3 nuclear locations; pixel intensities for each region were averaged and then used to calculate a nuclear:cytoplasmic ratio (ratio_nuc/cyt_). For each condition and replicate, individual cell ratios for at least 20 separate cells were used to calculate the average and standard deviation values displayed in figures. Results are reported as a ratio of the nuclear fluorescence to the cytoplasmic fluorescence (ratio_nuc/cyt_). Three independent, biological replicates were performed for each strain and condition tested.

### Metabolomics and evaluation of AMP, ADP, and ATP levels

DBY12000, cDBY0037, and cDBY0045 were grown overnight at 30°C in SD, subcultured into 50 mL of SD, and then grown with shaking at 30°C until reaching log phase growth with a cell concentration of approximately OD_600_= 0.6. Next, 1.25 OD_600_ units were collected onto a 0.45μm nylon membrane (Millipore HNWP02500), and the membrane was immediately quenched in 3mL -80°C extraction solvent (4:1 methanol:water, containing isotopically-labeled internal standards). The remaining culture was centrifuged, resuspended in 45mL of prewarmed SGal, and then returned to the incubator for 30 minutes. Another 1.25 OD_600_ units were collected and quenched following the same filtering procedure described above. Samples were collected from three independent, biological replicates. The membranes containing the cells were chilled in the extraction solvent for at least 20 minutes at -20°C. Each membrane was washed to collect all cellular material in the solvent, and the filters were discarded. Samples were centrifuged at 4°C for 10 mins at 3220 x *g*. The supernatant was collected and stored at -80°C for metabolomic analysis.

Metabolomics extracts were analyzed on an Agilent 1290 Bio HPLC coupled to a Sciex 7500 triple quadrupole mass spectrometer (MS). Chromatographic separation was achieved on a SeQuant® ZIC®-pHILIC 5µm 200Å 150 x 2.1mm HPLC column. Mobile phase A was 20 mM ammonium carbonate brought to pH 9.2 using ammonium hydroxide, with 1 µM medronic acid. Mobile phase B was 95% acetonitrile, 5% mobile phase A. The gradient was as follows at 150 µL/min flowrate: 0 min: 84.2 %B, 2.5 min: 76.9 %B, 5 min: 68.4 %B, 7.5 min: 60 %B, 10 min: 52.6 %B, 15 min: 36.8 %B, 20 min: 21.1 %B, 22 min: 15.8 %B, 22.5 min: 84.2 %B, 30 min: 84.2 %B. MS acquisition occurred in scheduled multiple reaction monitoring (MRM) mode with 2 transitions per compound whenever possible. Dwell time was set to a minimum of 4 ms, and a maximum of 250 ms, with a target cycle time of 1 second. Source conditions were as follows: ion source gas 1 60 psi, ion source gas 2 60 psi, curtain gas 46 psi, CAD gas 8, source temperature 350°C. Polarity switching was utilized with a 1600V spray voltage for positive mode, 1900V for negative, and a pause time of 4ms.

Data files were converted to mzML format using ProteoWizard msConvert (Chambers 2012), and all peaks were manually reviewed using MAVEN2 software (Seitzer 2022). All reported compounds were matched by retention time and a secondary transition whenever possible. Intensities were reported as the log2 value of smoothed peak area (including AMP, ADP, and ATP). For peaks with long tails where the full peak width exceeded the length of the MRM window (Uridine triphosphate, Fructose-1,6-bisphosphate, 1-Methylhistidine, Orotate) the topmost 3 points of each smoothed peak were instead used for quantification. Any peak below 1.5-fold the signal for the corresponding blanks’ median intensity was removed from the data set.

LCMS-grade acetonitrile, methanol and water were purchased from Fisher Scientific. LCMS-grade ammonium carbonate and ammonium hydroxide were purchased from Sigma. Isotopically labeled internal standards including 13C6-Fructose 6-Phosphate (p/n CLM-8962), 13C6-Glucose 6-Phosphate (p/n CLM-8367), 13C-Lactate (p/n CLM-1579), D4-Citrate (p/n DLM-3487), 13C4-Fumarate (p/n CLM-1529), d5-Phenylalanine (p/n DLM-1258), d4-Lysine (p/n DLM-2640), d4-Succinate (DLM-2307), 13C5-Glutamate (p/n CNLM-554-H) were purchased from Caymen Isotope Laboratories. Medronic acid was purchased from Agilent (p/n 5191-4506).

## ACKNOWLEDGEMENTS

We would like to thank Scott Emr for generously allowing us to use his laboratory facilities and fluorescent microscope. We would also like to thank Dan Gottschling for early conversations about this research and constructive comments regarding this manuscript. We would also like to thank the Cornell University Biotechnology Resource Center Genomics Facility (RRID:SCR_021727) for Sanger sequencing services to confirm plasmid constructs.

## Abbreviations

FGD: Failure of Glucose Derepression
ETC: electron transport chain

