## Supplemental Information for "Electron transport chain disruption leads to failure of glucose derepression via the Snf1/AMPK pathway"

**Supplemental Table 1.** Strains used in this study.

**Supplemental Figure 1.** Respiratory incompetent rho$^0$ cells display FGD.

**Supplemental Figure 2.** Observation of FGD in *coq2Δ*$_{gal}$ .

**Supplemental Figure 3.** FGD phenomenon is observed only under minimal media conditions.

**Supplemental Figure 4.** Average ratio$_{nuc/cyt}$ of biological replicates 2 and 3 for Mig1 localization during FGD.

**Supplemental Figure 5.** Tracking Mig1 localization in glucose repressed galactose grown strains.

**Supplemental Figure 6.** Average ratio$_{nuc/cyt}$ of biological replicates 2 and 3 for Snf1 localization during FGD.

**Supplemental Figure 7.** Retrograde signaling is not an intermediate pathway by which the ETC communicates with Snf1 under low glucose conditions.

**Supplemental Figure 8.** Decreased ADP levels do not correlate with FGD.

**Supplemental Figure 9.** Reduced Cat8 transcription factor activity observed in ETC mutants post-diauxic shift upon starvation in SD.

| Strain designation | Name in figures | Genotype/Description | Shown in figure | Ref |
|---|---|---|---|---|
| DBY12000 | Wild type, WT | S288C/FY derivative. MAT**a** prototroph. $HAP1^+$ | 1, 3, SF1, SF2, SF3, SF5, SF6, SF7 | See Materials and Methods |
| PGY613 | $coq2\Delta_{gal}$ | MAT**a** $HAP1^+$ *coq2*Δ::kanMX (maintained only on galactose) | SF2, SF3 | This study |
| PGY614 | $cox4\Delta_{gal}$ | MAT**a** $HAP1^+$ *cox4*Δ::kanMX (maintained only on galactose) | 1, 3, SF3, SF5 | This study |
| cDBY0083 | mt NADH dehydrogenase mutant | MAT**a** $HAP1^+$ *ndi1*Δ::kanMX *nde1*Δ::hphMX *nde2*Δ::bleMX | SF7 | Lewis A, 2021 |
| cDBY0037 | $coq2\Delta_{glu}$ | MAT**a** $HAP1^+$ *coq2*Δ::kanMX | SF2, SF6 | Lewis A, 2021 |
| cDBY0045 | $cox4\Delta_{glu}$, Complex IV mutant | MAT**a** $HAP1^+$ *cox4*Δ::kanMX | 1, SF6, SF7 | Lewis A, 2021 |
| PGY600 | $ura3\Delta_{gal}$ | MAT**a** $HAP1^+$ *ura3*Δ0 (maintained only on galactose) | 1 | This study |
| PGY604 | $cox4\Delta ura3\Delta_{gal}$ | MAT**a** $HAP1^+$ *cox4*Δ*ura3*Δ0 (maintained only on galactose) | 1 | This study |
| DBY12007 | S288C | S288C/FY derivative. MAT**a**/α $HAP1^+$ prototrophic diploid | 2 | See Materials and Methods |
| DBY15119 | W303 | W303 diploid prototroph | 2 | Ralser M, 2012 |
| PGY4 | Simi White | Commercial wine strain[a] | 2 | Richter CL, 2013 |
| PGY7 | CSM | Commercial wine strain[a] | 2 | Richter CL, 2013 |
| PGY34 | YPS1000 | Natural isolate from New Jersey, United States; from oak exudate[b] | 2 | Sniegowski PD, 2002 |
| PGY46 | Bb32 | Natural isolate from Ravenswood Winery, California, United States; isolated by Robert Mortimer from a Zinfandel in 1993; progenitor of RM11-1a[c] | 2 | Mortimer RK, 1994 |
| PGY664 | S288C $\rho^0$ | DBY12007 diploid $\rho^0$ (maintained only on galactose) | 2 | This study |
| PGY665 | W303 $\rho^0$ | W303 diploid $\rho^0$ (maintained only on galactose) | 2 | This study |
| PGY666 | Simi White $\rho^0$ | Simi White $\rho^0$ (maintained only on galactose) | 2 | This study |
| PGY667 | CSM $\rho^0$ | CSM $\rho^0$ (maintained only on galactose) | 2 | This study |
| PGY668 | YPS1000 $\rho^0$ | YPS1000 $\rho^0$ (maintained only on galactose) | 2 | This study |
| PGY669 | Bb32 $\rho^0$ | Bb32 $\rho^0$ (maintained only on galactose) | 2 | This study |
| DBY12785 | *mig1Δ* | MATa $HAP1^+$ *mig1*Δ::hphMX | 3 | This study |
| PGY731 | $mig1\Delta coq2\Delta_{gal}$ | MAT**a** $HAP1^+$ *mig1*Δ::hphMX *coq2*Δ::kanMX (maintained only on galactose) | 3 | This study |
| PGY733 | $mig1\Delta cox4\Delta_{gal}$ | MAT**a** $HAP1^+$ *mig1*Δ::hphMX *cox4*Δ::kanMX (maintained only on galactose) | 3 | This study |
| DBY12509 | *hxk2Δ* | MATa $HAP1^+$ *hxk2*Δ::bleMX | 3 | This study |
| PGY735 | $hxk2\Delta coq2\Delta_{gal}$ | MAT**a** $HAP1^+$ *hxk2*Δ::bleMX *coq2*Δ::kanMX (maintained only on galactose) | 3 | This study |
| PGY736 | $hxk2\Delta cox4\Delta_{gal}$ | MAT**a** $HAP1^+$ *hxk2*Δ::bleMX *cox4*Δ::kanMX (maintained only on galactose) | 3 | This study |
| PGY720 | Wild type | MAT**a** $HAP1^+$ *mig1*::*MIG1*-mNeonGreen-Ca*URA3* *hmg1*::*HMG1*-yomRuby2-kanMX (maintained only on galactose) | 4, SF4, SF8 | This study |
| PGY722 | $coq2\Delta_{gal}$ | MAT**a** $HAP1^+$ *mig1*::*MIG1*-mNeonGreen-Ca*URA3* *hmg1*::*HMG1*-yomRuby2-kanMX *coq2*Δ::kanMX (maintained only on galactose) | 4 | This study |
| PGY723 | $cox4\Delta_{gal}$ | MAT**a** $HAP1^+$ *mig1*::*MIG1*-mNeonGreen-Ca*URA3* *hmg1*::*HMG1*-yomRuby2-kanMX *cox4*Δ::kanMX (maintained only on galactose) | 4, SF4, SF8 | This study |
| PGY725 | Wild type | MAT**a** $HAP1^+$ *snf1*::*SNF1*-mNeonGreen-natMX *hmg1*::*HMG1*-yomRuby2-kanMX (maintained only on galactose) | 5 | This study |
| PGY727 | $coq2\Delta_{gal}$ | MAT**a** $HAP1^+$ *snf1*::*SNF1*-mNeonGreen-natMX *hmg1*::*HMG1*-yomRuby2-kanMX *coq2*Δ::kanMX (maintained only on galactose) | 5 | This study |
| PGY729 | $cox4\Delta_{gal}$ | MAT**a** $HAP1^+$ *snf1*::*SNF1*-mNeonGreen-natMX *hmg1*::*HMG1*-yomRuby2-kanMX *cox4*Δ::kanMX (maintained only on galactose) | 5 | This study |
| PGY464 | *rtg3*Δ | MATa $HAP1^+$ *rtg3*Δ::kanMX | SF5 | This study |

**Supplemental Table 1. Strains used in this study.**

a - Kind gift from Tom Pugh and E&J Gallo Winery (Modesto, CA).
b - Kind gift from Joseph Schacherer Lab.
c - Kind gift from Leonid Kruglyak Lab.

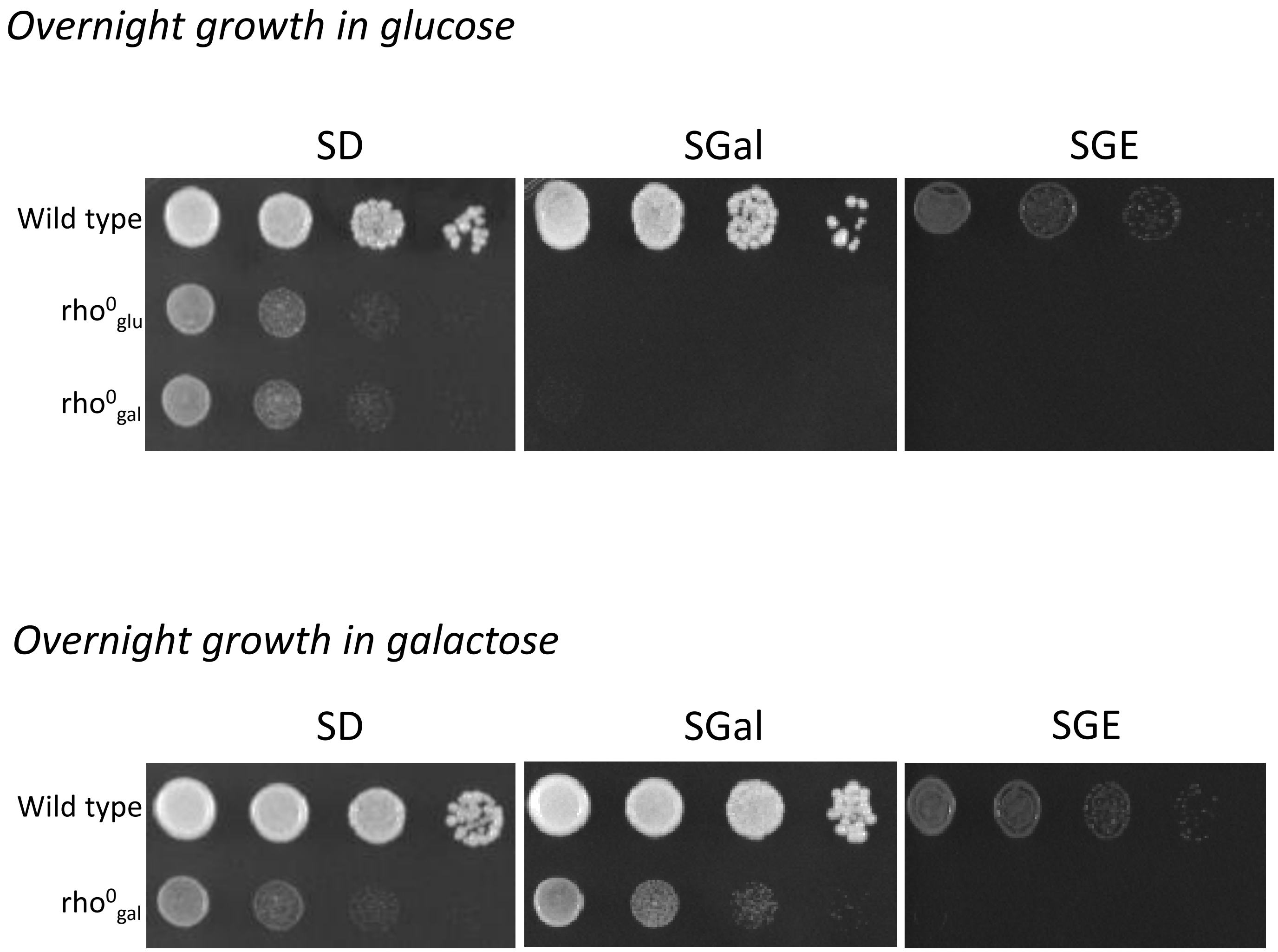


**Supplemental Figure 1. Respiratory incompetent rho$^0$ cells display FGD.** Wild type (WT; DBY12000), rho$^0_{glu}$(DBY12000 treated with EtBr; generated and maintained only in glucose), rho$^0_{gal}$(DBY12000 treated with EtBr; generated and maintained only in galactose) were grown overnight in either minimal glucose (SD) or minimal galactose (SGal). The following day, 1:10 dilutions were prepared, and cultures were pronged onto SD, SGal and SGE. Plates were incubated at 30℃ and photographed after 3 days. Shown are representative images from at least 4 biological replicates.

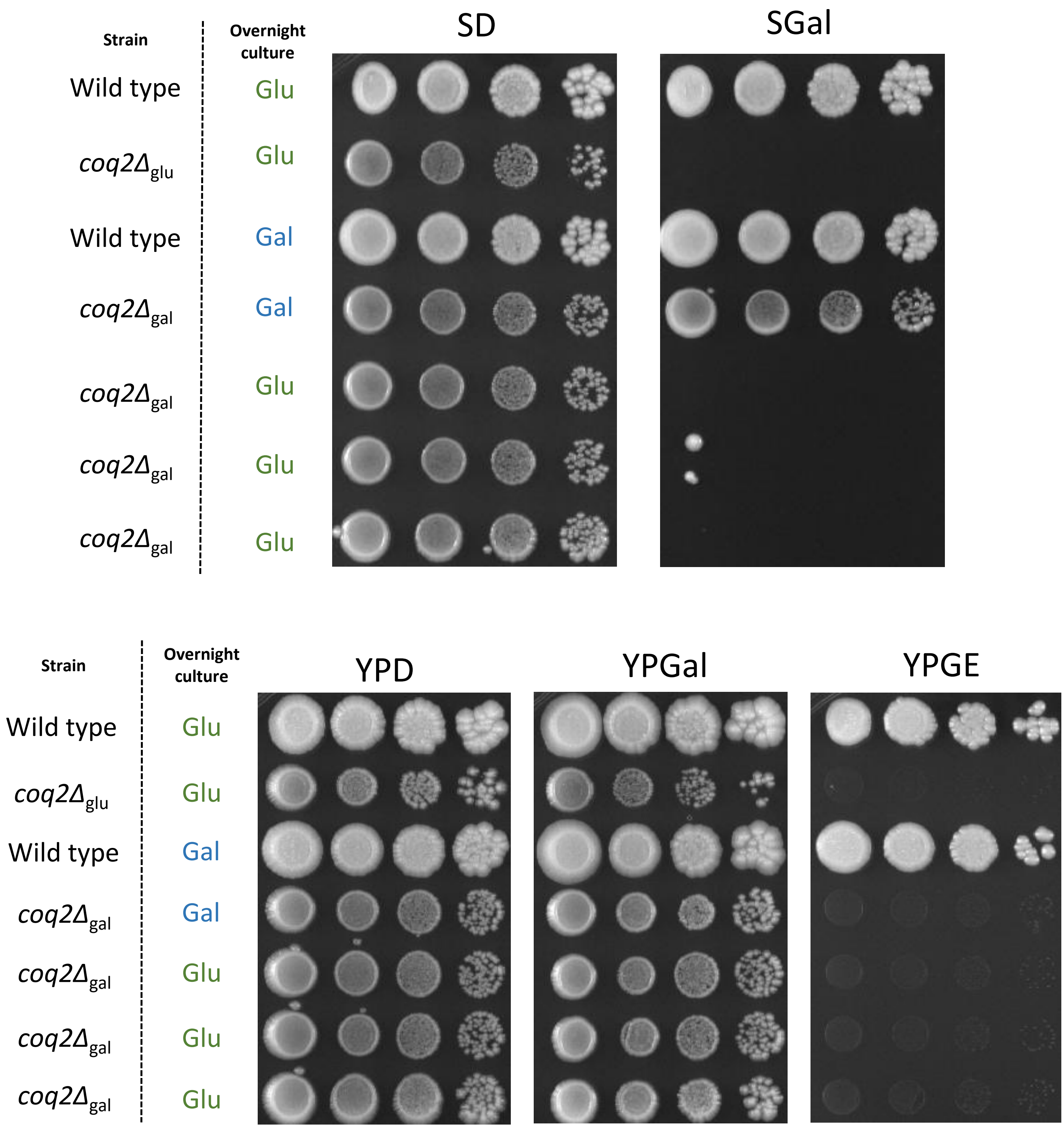


**Supplemental Figure 2. Observation of FGD in *coq2Δ*$_{gal}$.** Wild type (DBY12000), *coq2Δ*$_{glu}$, *coq2Δ*$_{gal}$ were grown overnight in either SD or SGal. Strains were washed the next day 3X in sterile MilliQ water, 10-fold dilutions were prepared, and cells were pronged onto SD, SGal, YPD, YPGal and YPGE. Plates were allowed to incubate at 30°C for 6 days before photographing.

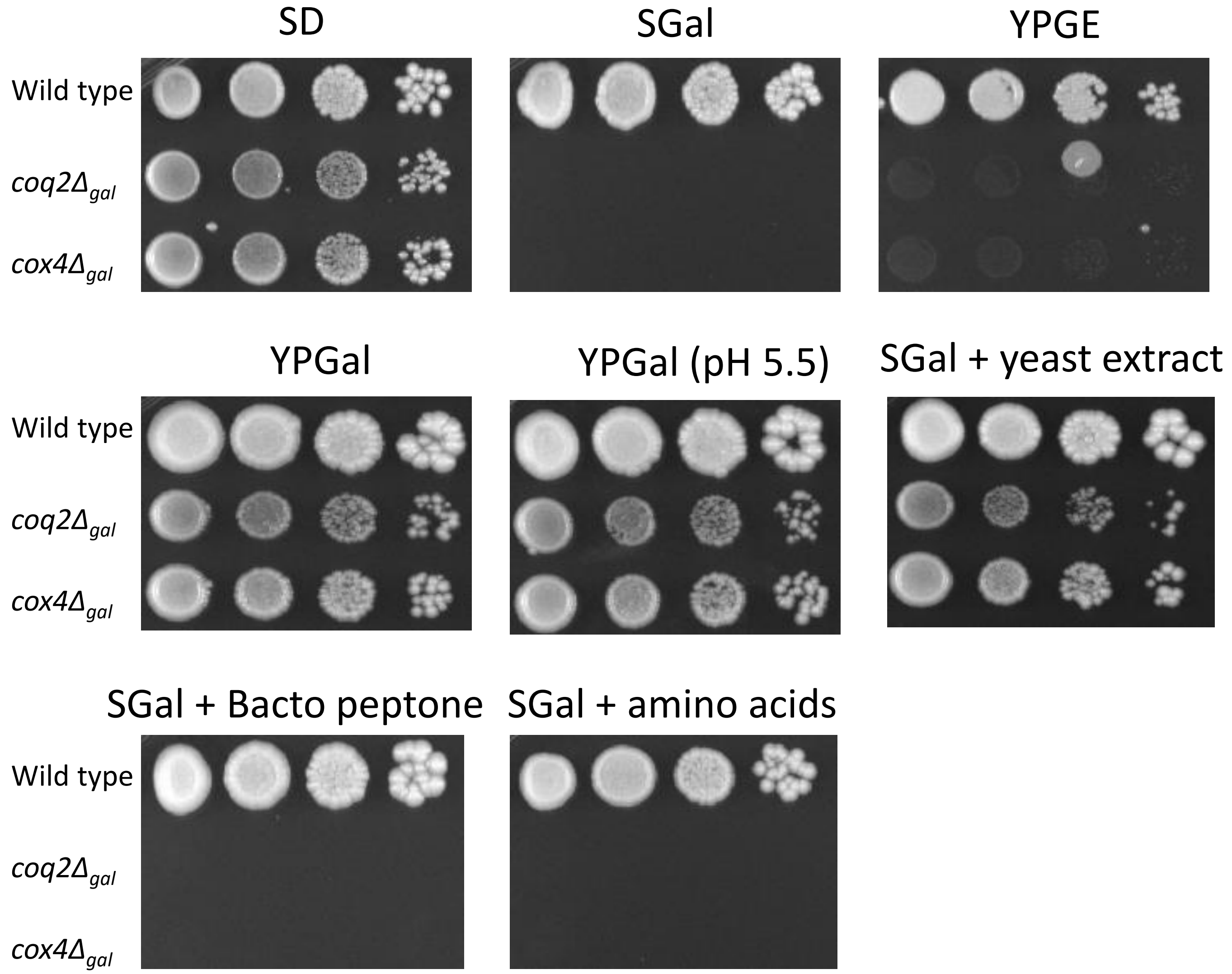


**Supplemental Figure 3**. **FGD phenotype is observed only under minimal media conditions.** Indicated strains were grown overnight in SD. The next day, cultures were washed, cells collected, 1:10 dilutions were prepared and pronged onto SD, SGal, YPGE, YPGal, YPGal (pH 5.5), SGal+yeast extract, SGal+Bacto peptone and SGal+amino acid mixture from SC-supplemented mix. Plates were incubated at 30°C and photographed after 4 days.

**A.**

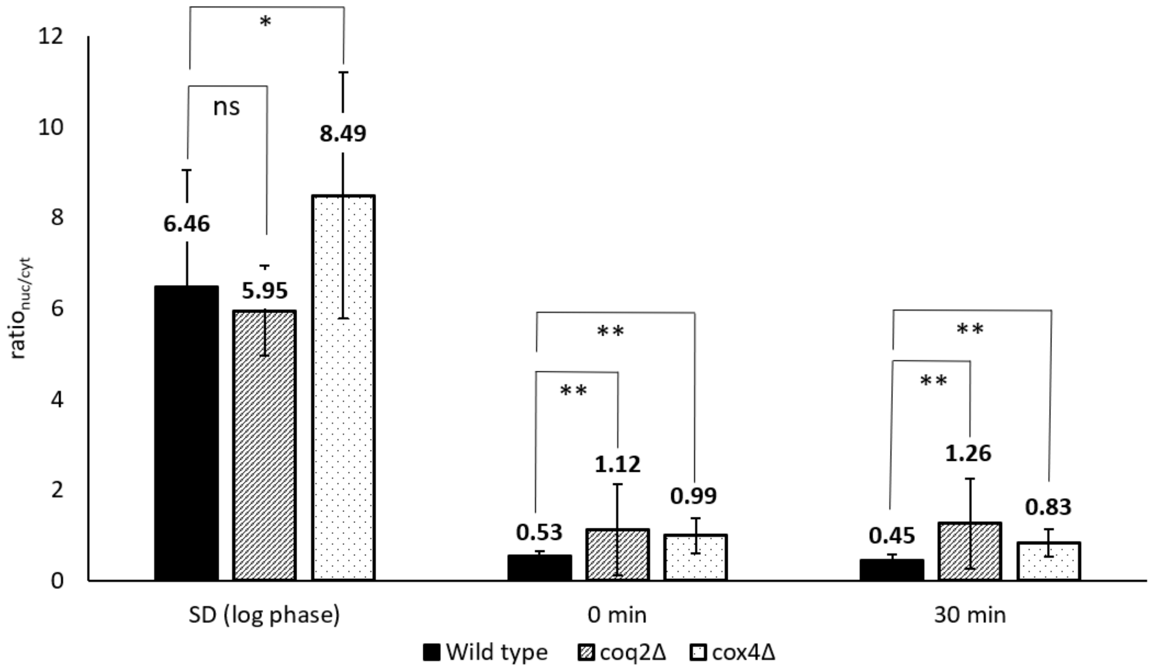


**B.**

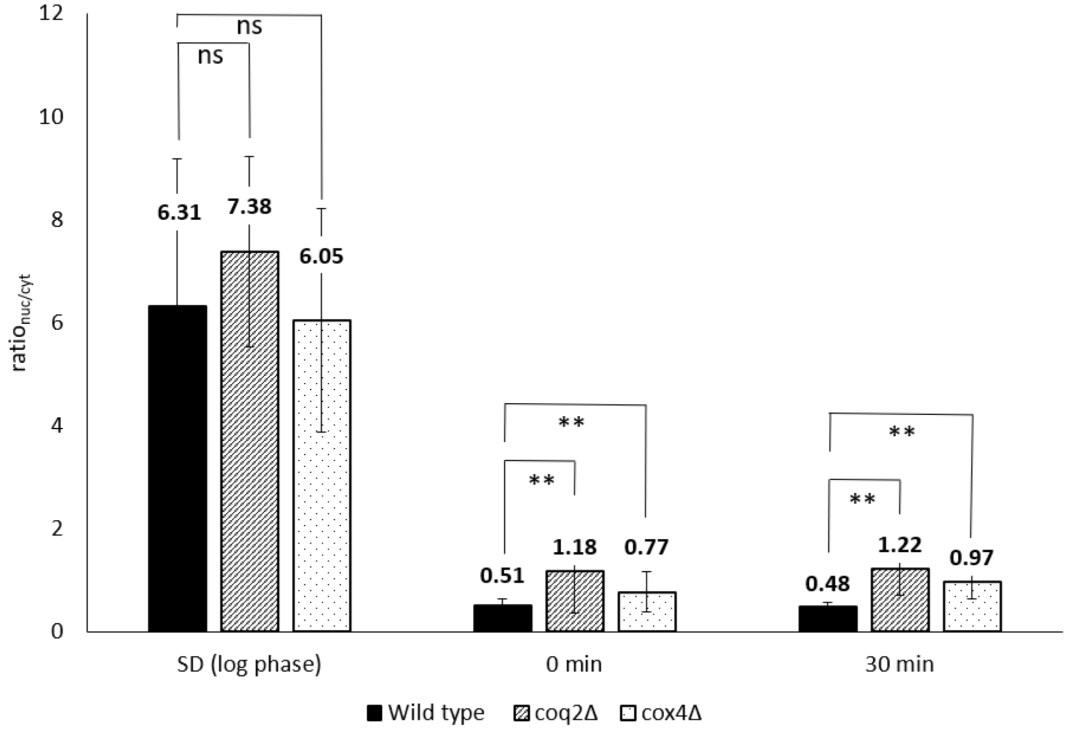


**Supplemental Figure 4**. **Average ratio$_{nuc/cyt}$ of biological replicates 2 and 3 for Mig1 localization during FGD.** Bar graphs of average ratio$_{nuc/cyt}$ of replicate 2 (A) and replicate 3 (B) of Mig1 localization studies in galactose-maintained wild type**,** *coq2Δ*$_{gal}$ and *cox4Δ* $_{gal}$ strains during FGD (quantification of the first replicate is shown in Figure 4D). Cells were grown in SGal overnight before being transferred to SD. Once the culture had reached exponential growth phase in SD, cells were collected by centrifugation, washed with sterile milli-Q water and subcultured into fresh SGal at an initial $OD_{600}$ of 0.2. Images were taken at indicated time points ('0h' cells were collected immediately after resuspension into fresh SGal), and 20-30 cells were visualized and quantified per replicate. *: uncorrected *p* value < 0.05, **: uncorrected *p* value < 0.001, ns: not significant ($p \geq 0.05$). For each strain, at least 20 cells per time point were counted to calculate ratio$_{nuc/cyt}$ from a single replicate.

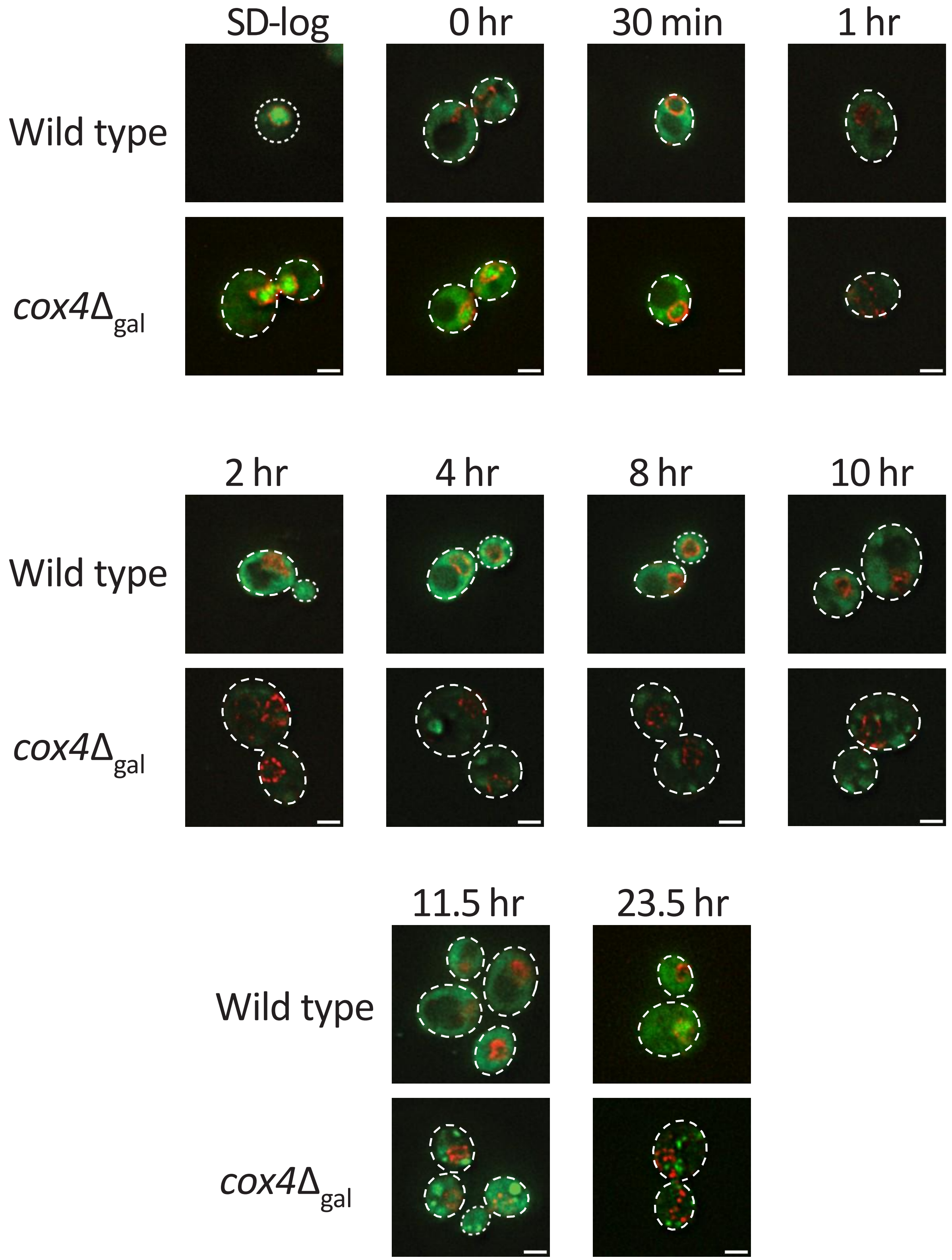


**Supplemental Figure 5**. **Tracking Mig1 localization in glucose repressed galactose grown strains.** Indicated strains were maintained and grown on minimal galactose medium before cells were subcultured into SD. After cells had attained exponential growth phase in SD ($OD_{600}$ = 0.2), cells were washed and transferred to SGal medium. At stated time points, cells were imaged for Mig1 localization using the DeltaVision Elite system. Shown are merged images of mNeonGreen (for Mig1) and yomRuby2 (for Hmg1) filters. Three replicates were performed with 20-30 cells visualized per replicate. All images shown are representative of one replicate. Scale bar = 2µm.

**A.**

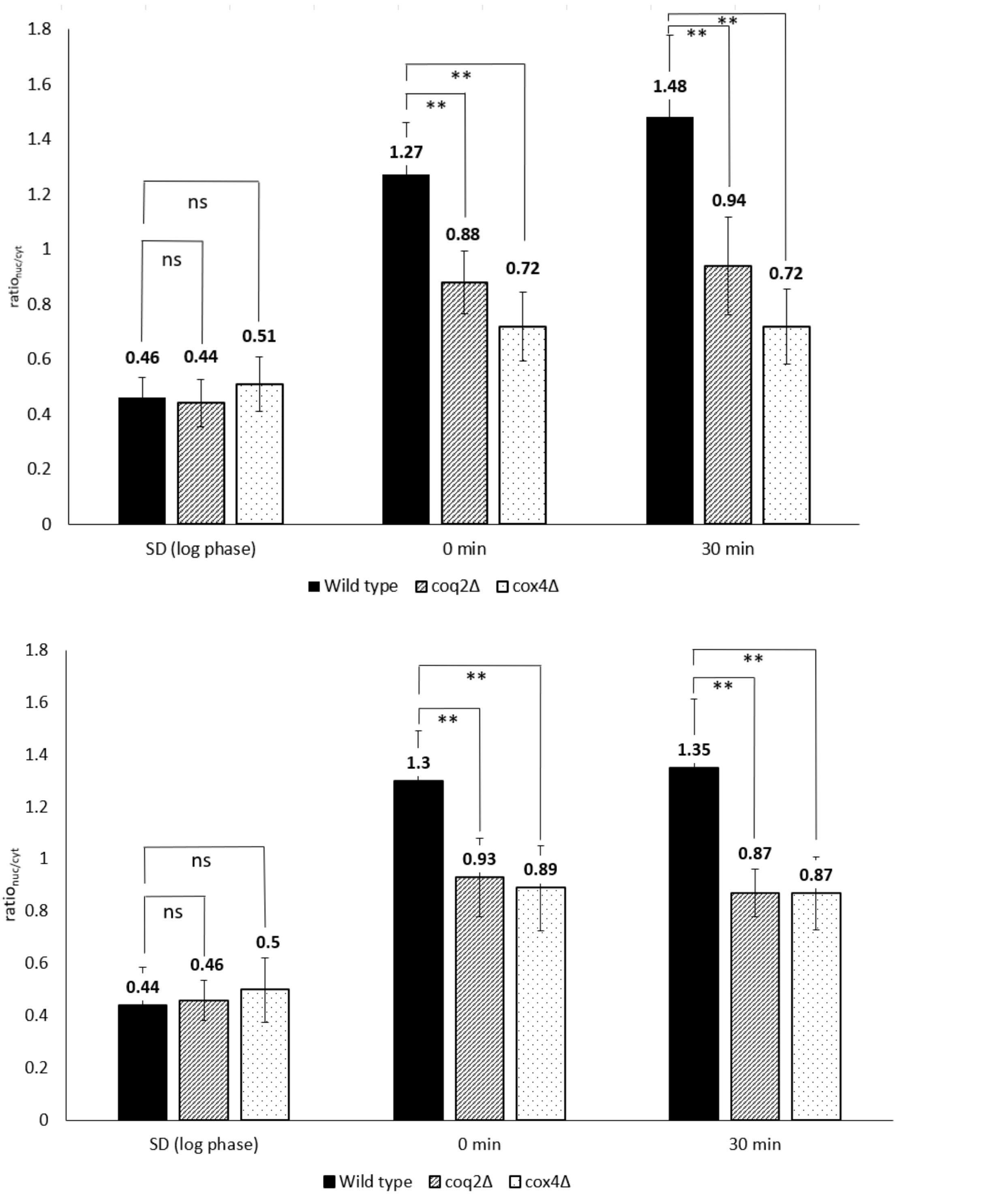


**B.**



**Supplemental Figure 6**. **Average ratio$_{nuc/cyt}$ of biological replicates 2 and 3 for Snf1 localization during FGD.** Bar graphs of average ratio$_{nuc/cyt}$ of replicate 2 (A) and replicate 3 (B) of Snf1 localization studies in galactose-maintained wild type**,** *coq2Δ*$_{gal}$ and *cox4Δ* $_{gal}$ strains during FGD (quantification of the first replicate is shown in Figure 5D). Cells were grown in SGal overnight before being transferred to SD. Once the culture had reached exponential growth phase in SD, cells were collected by centrifugation, washed with sterile milli-Q water and subcultured into fresh SGal at an initial $OD_{600}$ of 0.2. Images were taken at indicated time points ('0h' cells were collected immediately after resuspension into fresh SGal), and 20-30 cells were visualized and quantified per replicate. *: uncorrected *p* value < 0.05, **: uncorrected *p* value < 0.0001, ns: not significant ($p \geq 0.05$). For each strain, at least 20 cells per time point were counted to calculate ratio$_{nuc/cyt}$ from a single replicate.

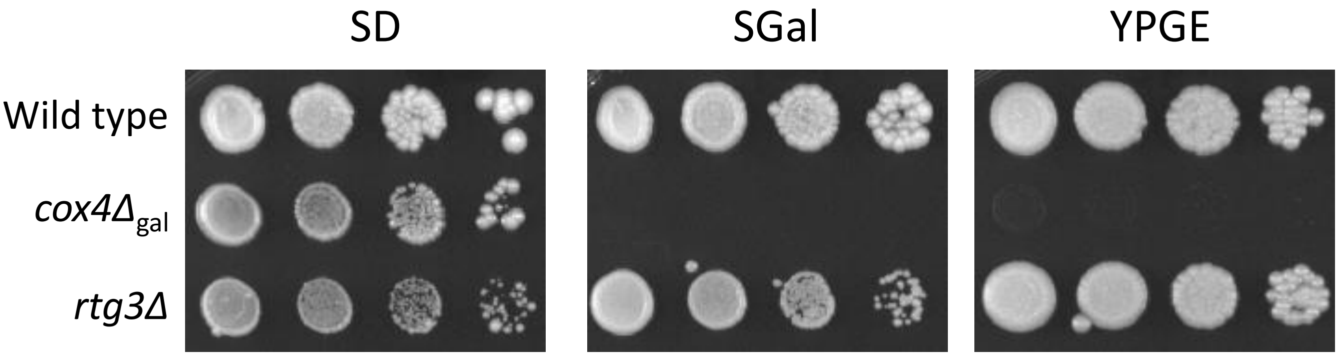


**Supplemental Figure 7. Retrograde signaling is not an intermediate pathway by which the ETC communicates with Snf1 under low glucose conditions.** Indicated strains were grown overnight in SD before cells were washed, collected by centrifugation, 10-fold serial dilutions prepared and pronged onto SD, SGal and YPGE. Plates were incubated at 30°C for 7 days before photographing. Shown are representative images of at least three biological replicates.

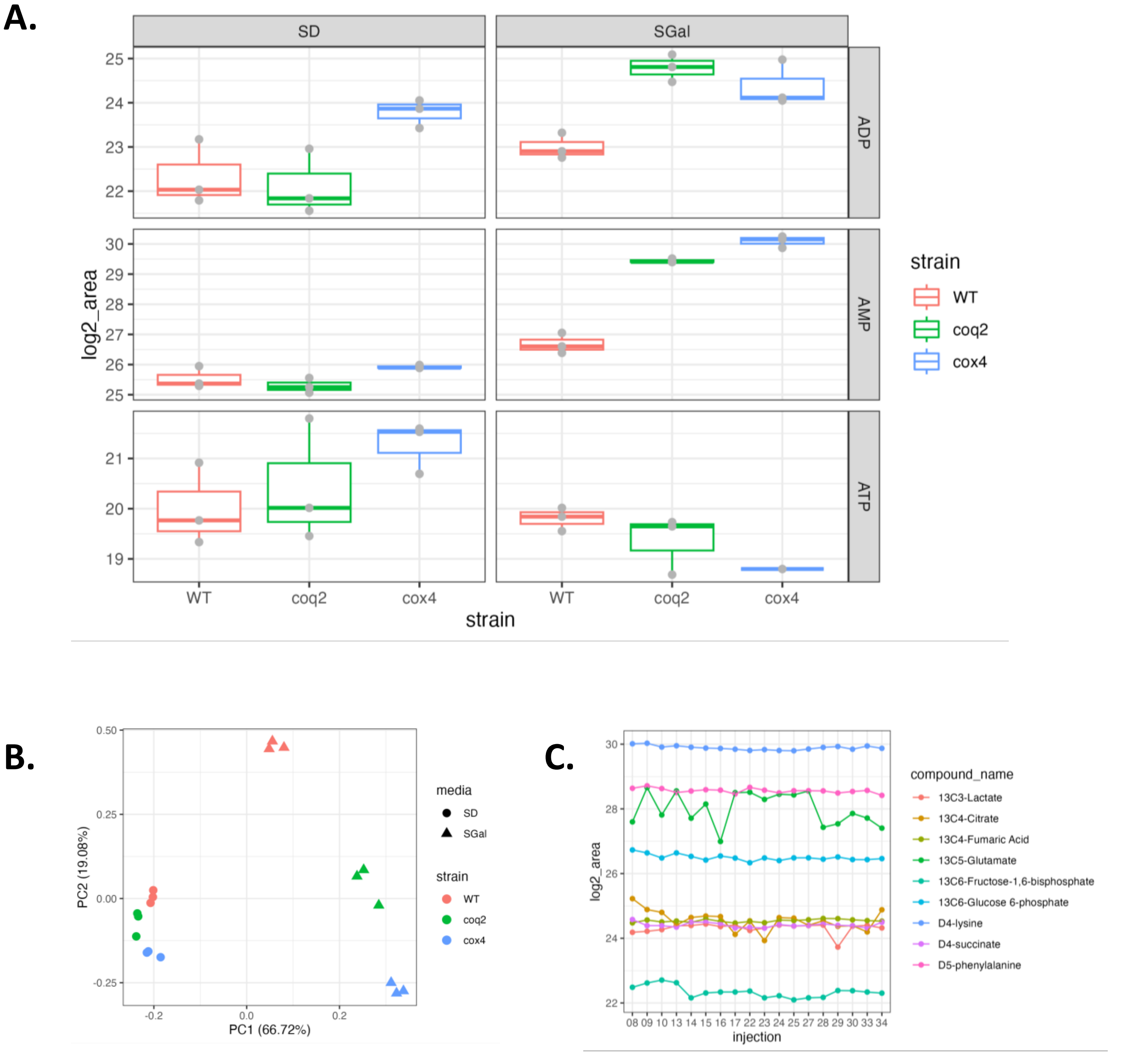


**Supplemental Figure 8. Decreased ADP levels do not correlate with FGD. A.** Relative AMP, ADP and ATP levels of ETC mutants (*coq2Δ* and *cox4Δ*) compared to wild type during log phase growth in SD and after a 30-minute incubation in SGal. Values represent smoothed area under the curves for the indicated metabolites. Data are derived from 3 independent biological replicates. **B**. Principal component analysis of high-quality metabolites from each sample illustrates replicates cluster close together, glucose-grown strains cluster together, but ETC mutants and wild type do not cluster together after galactose incubation. **C**. Internal standard peak intensity across run time. **D.** Heat-map of high-quality metabolites identified for each strain, condition, and replicate during metabolomics experiment. Values represent the $\log_2$ smoothed peak area relative to the median of that compound across all samples.

**D.**

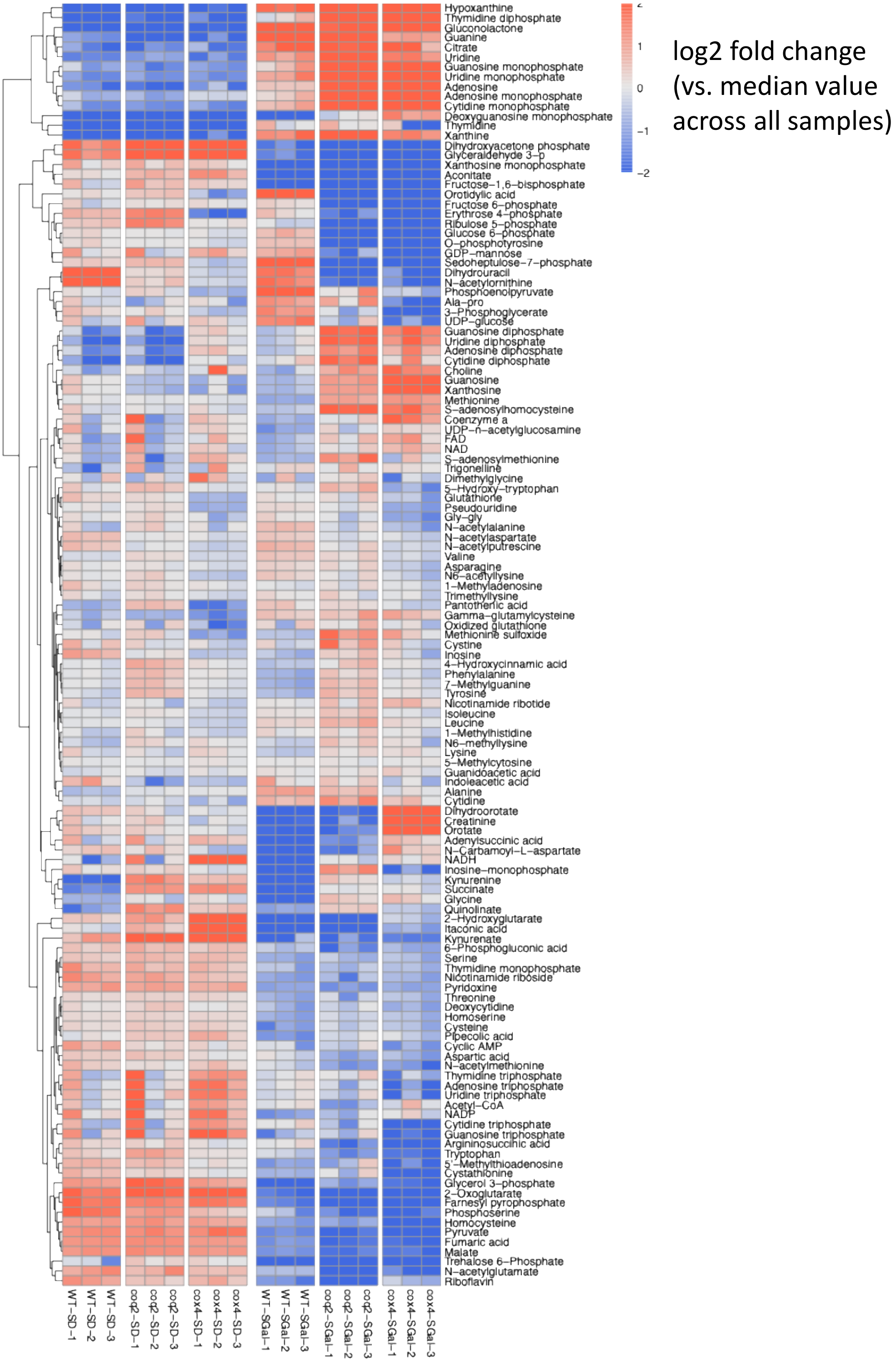


**Supplemental Figure 8. Decreased ADP levels do not correlate with FGD. A.** Relative AMP, ADP and ATP levels of ETC mutants (*coq2Δ* and *cox4Δ*) compared to wild type during log phase growth in SD and after a 30-minute incubation in SGal. Values represent smoothed area under the curves for the indicated metabolites. Data are derived from 3 independent biological replicates. **B**. Principal component analysis of high-quality metabolites from each sample illustrates replicates cluster close together, glucose-grown strains cluster together, but ETC mutants and wild type do not cluster together after galactose incubation. **C**. Internal standard peak intensity across run time. **D.** Heat-map of high-quality metabolites identified for each strain, condition, and replicate during metabolomics experiment. Values represent the $\log_2$ smoothed peak area relative to the median of that compound across all samples.

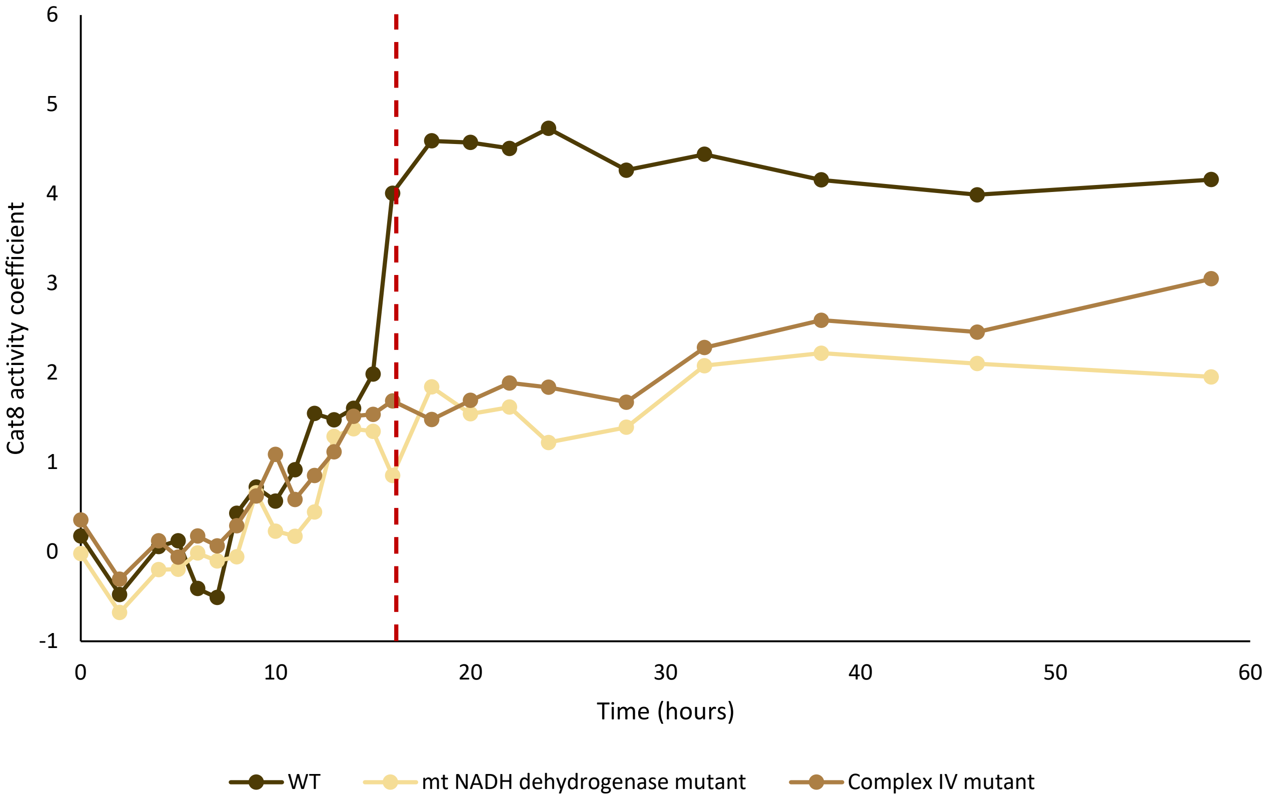


**Supplemental Figure 9. Reduced Cat8 transcription factor activity observed in ETC mutants post-diauxic shift upon starvation in SD.** Activity of Cat8 in wild type (WT) and ETC mutants (mt NADH dehydrogenase [*ndi1*Δ*nde1*Δ*nde2*Δ] and Complex IV [*cox4*Δ] mutants). Post-diauxic shift (around 16h and indicated by red dotted line), the activity of Cat8 is downregulated in the ETC mutants as compared to wild type. Image reconstructed from data downloaded from *Lewis et al.,* 2021.
